# YAP antagonizes TEAD-mediated AR signaling and prostate cancer growth

**DOI:** 10.1101/2022.08.01.502314

**Authors:** Shu Zhuo, Xu Li, Yong Suk Cho, Yuchen Liu, Yingzi Yang, Jian Zhu, Jin Jiang

## Abstract

Hippo signaling restricts tumor growth by inhibiting the oncogenic potential of YAP/TAZ-TEAD transcriptional complex. Here we uncover a context-dependent tumor suppressor function of YAP in androgen receptor (AR) positive prostate cancer (PCa) and show that YAP impedes AR^+^ PCa growth by antagonizing TEAD-mediated AR signaling. TEAD forms a complex with AR to enhance its promoter/enhancer occupancy and transcriptional activity. YAP and AR compete for TEAD binding and consequently, elevated YAP in the nucleus disrupts AR-TEAD interaction and prevents TEAD from promoting AR signaling. Pharmacological inhibition of Hippo/MST1/2 kinase or transgenic activation of YAP suppressed the growth of PCa expressing therapy resistant AR splicing variants. Our study uncovers an unanticipated crosstalk between Hippo and AR signaling pathways, reveals an antagonistic relationship between YAP and TEAD in AR^+^ PCa, and suggests that targeting the Hippo signaling pathway may provide a therapeutical opportunity to treat PCa driven by therapy resistant AR variants.

- YAP acts as a context-dependent tumor suppressor in AR^+^ PCa
- TEAD interacts with AR to enhance its promoter/enhancer occupancy
- YAP inhibits AR activity by competing for TEAD binding
- Small molecule Hippo pathway inhibitor impedes AR^+^ PCa growth

## Introduction

Prostate cancer (PCa) is the second most commonly diagnosed malignancy in USA, where approximately one in every seven men will be diagnosed with the disease during their life span (Siegel *et al*, 2018). The majority of diagnosed PCa are androgen receptor (AR) positive with AR as the main driver of cancer progression (Dai *et al*, 2017). Although anti-hormone therapy such as androgen deprivation and androgen receptor (AR) inhibition can effectively control AR^+^ PCa progression, many treated patients will eventually develop resistance and give rise to castration-resistant prostate cancer (CRPC)(Sharifi *et al*, 2005). Hence, it is necessary to develop new therapeutic strategies to overcome therapy resistance.

The Hippo tumor suppressor pathway is an evolutionarily conserved signaling pathway that controls tissue growth, organ size, tissue regeneration and cancer progression (Halder & Johnson, 2011; Pan, 2007; Yu *et al*, 2015; Zhang *et al*, 2009). In the core Hippo pathway, the upstream kinase Hippo (Hpo)/MST1/2 phosphorylates and activates the downstream kinase LATS1/2 (Harvey *et al*, 2003; Jia *et al*, 2003; Pantalacci *et al*, 2003; Udan *et al*, 2003; Wu *et al*, 2003), leading to phosphorylation and inactivation of the transcriptional effector YAP/TAZ (Dong *et al*, 2007; Lei *et al*, 2008; Zhao *et al*, 2007). Inhibition of Hippo signaling allows YAP/TAZ to translocate into the nucleus where it binds the Hippo pathway transcription factor TEAD to regulate target gene expression (Goulev *et al*, 2008; Wu *et al*, 2008; Zhang *et al*, 2008; Zhao *et al*, 2008). *YAP* is amplified and its protein level and nuclear localization are elevated in many types of human cancer including liver, lung, colon, and ovarian cancers (Yu *et al*., 2015; Zanconato *et al*, 2016). Furthermore, YAP overexpression and knockout of MST1/2 or other upstream components of the Hippo pathway in mouse liver resulted in hepatomegaly, leading to hepatocellular carcinoma formation (Camargo *et al*, 2007; Dong *et al*., 2007; Lee *et al*, 2010; Lu *et al*, 2010; Song *et al*, 2010; Zhang *et al*, 2010; Zhou *et al*, 2009). These observations lead to a prevalent view that Hippo signaling functions as a tumor suppressor pathway by blocking the oncogenic potential of YAP/TAZ (Pobbati & Hong, 2020; Thompson, 2020; Yu *et al*., 2015; Zanconato *et al*., 2016). However, recent studies have revealed that YAP could function as tumor suppressor in several types of cancer although the mechanisms of action vary depending on the cancer types (Cheung *et al*, 2020; Cottini *et al*, 2014; Li *et al*, 2020; Ma *et al*, 2021; Moya *et al*, 2019).

Here, we investigated the role of Hippo signaling in AR positive PCa and explored the mechanism by which Hippo signaling acts through YAP to modulate AR signaling activity. We found that inhibition of Hippo signaling pathway or ectopic activation of YAP blocked the AR transcriptional program and AR^+^ PCa growth. We uncovered a previously uncharacterized function of TEAD in assisting AR signaling by forming a complex with AR to increase its promoter/enhancer occupancy. We demonstrated that YAP inhibits AR transcriptional activity and AR binding to its target promoters by disrupting the AR-TEAD signaling complex. We further showed that pharmacological inhibition of Hippo/MST1/2 impaired the growth of therapy resistant PCa *in vivo*. Our study thus unravels a nonconventional function of Hippo signaling in cancer and suggests that targeting the Hippo pathway could be a potential strategy to overcome endocrine therapy resistance in AR^+^ PCa patients.

## Result

### Activation of YAP suppresses AR^+^ PCa growth

Previous studies revealed a conflicting role of YAP in prostate cancer. While several studies indicated that YAP protein level was upregulated in PCa patient samples, which implied an oncogenic function of YAP (Dong *et al*., 2007; Zhao *et al*., 2007), others revealed that YAP was downregulated in PCa compared with normal prostate tissue (Collak *et al*, 2017; Hu *et al*, 2015). We speculated that YAP signaling could vary in PCa depending on AR signaling status. By analyzing TCGA data, we found that the expression levels of *YAP* and *TAZ* as well as their target gene *AMOTL2* were lower whereas the AR target genes *KLK2* and *KLK3* (also called PSA) were higher in 497 PCa samples compared with 52 normal tissues (Fig. EV1A). In AR activity high PCa samples, YAP signaling activity, which is represented by the expression levels of *YAP* and YAP signature target genes including *AMOTL2*, *CTGF* and *CYR61*, is negatively correlated with AR signaling activity, which is reflected by the expression levels of *KLK2* and *KLK3* (Fig. EV1B-D).

To determine the effect of YAP activation on AR^+^ PCa growth, we used the lentiviral system to ectopically express either wild type YAP (YAP-WT) or a constitutively active form of YAP (YAP-5SA) that lacks five LATS1/2 phosphorylation sites (Zhao *et al*., 2007) in AR^+^ PCa cell lines that expressed either full-length AR (LNCaP and C4-2) or AR splicing variants (22RV1 and R1-D567) (Horoszewicz *et al*, 1980; Nyquist *et al*, 2013; Sramkoski *et al*, 1999; Wu *et al*, 1994). Expression of YAP-5SA inhibited all four AR^+^ PCa cell growth (Fig. 1A-D). C4-2 and 22RV1 cells were relatively more sensitive to YAP activation as overexpression of YAP-WT also inhibited their proliferation albeit less effectively compared with YAP-5SA (Fig. 1B, C). Inactivation of Hippo signaling by treating cells with a small molecule compound XMU-MP-1, which specifically inhibits the MST1/2 kinase activity (Fan *et al*, 2016), also inhibited the growth of LNCaP, C4-2, 22RV1 and R1-D567 cancer cells (Fig. 1E-H). Immunostaining or fractionation of C4-2 cells treated with XMU-MP-1 confirmed that nuclear levels of YAP were increased by drug treatment (Fig. 1I-J). Knockdown of YAP by two independent siRNAs (siYAP_1 and siYAP_2) slightly increased C4-2 cell growth and reversed the inhibitory effect of XMU-MP-1(Fig. 1K-L), suggesting that XMU-MP-1 inhibited C4-2 growth at least in part through activation of YAP. Blocking Hippo signaling pathway by knockdown of both LATS1 and LATS2 (LATS1/2 KD) also inhibited the growth of LNCaP, C4-2, and 22RV1 cancer cells (Fig, 1M, N; Fig. EV2). Furthermore, the inhibition of C4-2 growth by LATS1/2 KD was reversed by YAP RNAi (Fig, 1O,P). Hence, activation of YAP either by transgenic overexpression or blockage of Hippo signaling pathway inhibits AR^+^ PCa growth *in vitro*.

**Figure 1.**
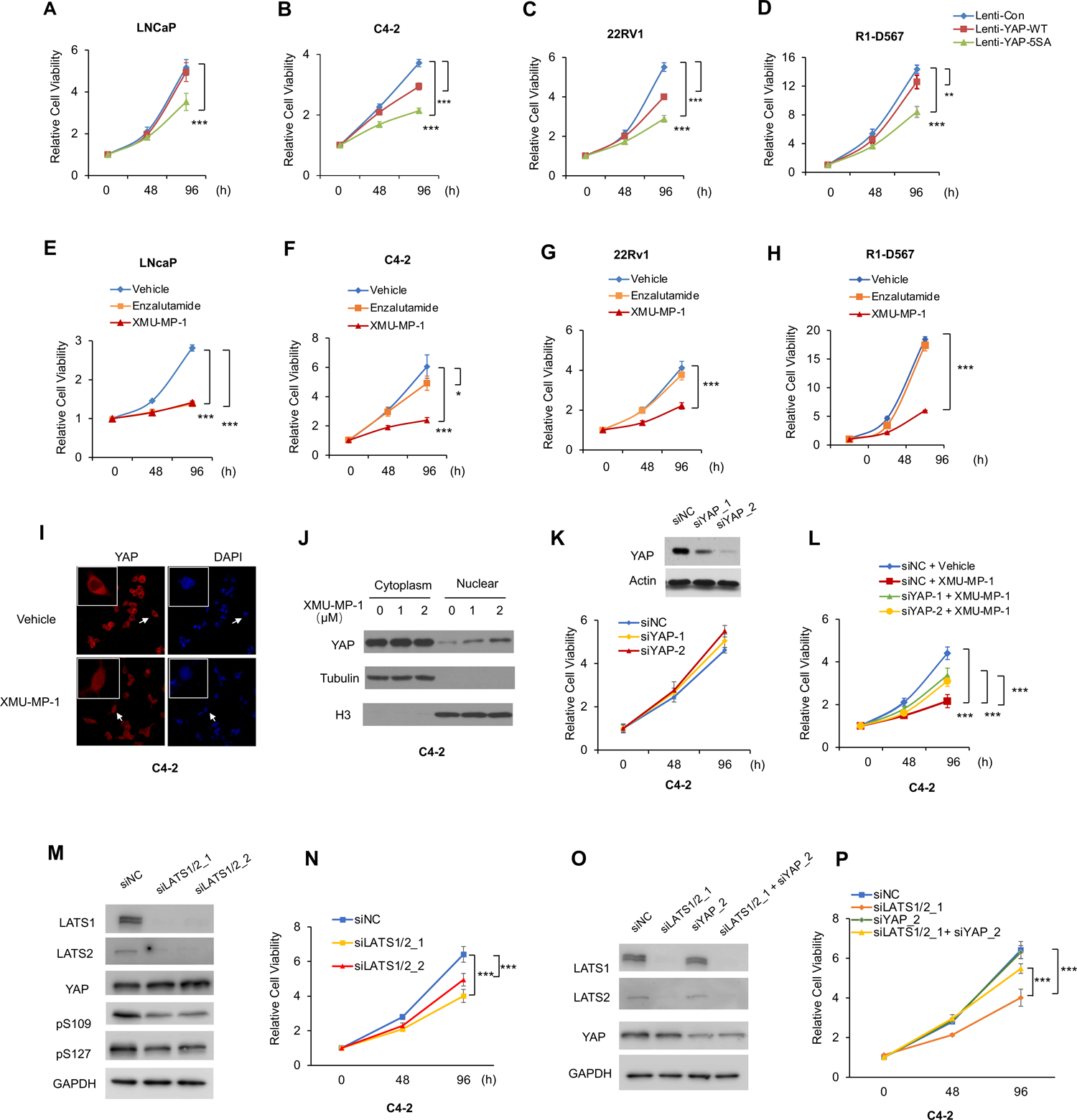
MST1/2 inhibition or YAP activation inhibits AR^+^ PCa growth *in vitro*. (A-D) Relative growth of the indicated prostate cancer cell lines expressing control, YAP-WT, or YAP-5SA constructs through lentiviral infection. (E-H) Relative growth of the indicated prostate cancer cell lines treated with XMU-MP-1 (1 μM for C4-2; 2 μM for LNCaP and 22RV1; 5 μM for R1-D567) or 5 μM Enzalutamide. (I) C4-2 cells treated with vehicle or 2 μM XMU-MP-1 were immunostained with DAPI anti-YAP antibody. Insets are enlarged views of cells indicated by the arrows. (J) Western blot analysis of YAP in cytoplasmic and nuclear fractions of C4-2 cells treated with XMU-MP-1 at the indicated concentrations. Tubulin and Histone 3(H3) mark the cytoplasmic and nuclear fractions, respectively. (K) Western blot analysis of YAP in (top) or relative growth of (bottom) C4-2 cells treated with two independent YAP siRNAs (siYAP_1 and siYAP_2) or control siRNA (siNC), (L) Relative growth of C4-2 cells treated with YAP or control siRNA and with vehicle or 1 μM XUM-MP-1. (M) Western blot analysis of LATS1, LATS2, YAP, phosphorylated YAP at S109 (pS109) and S127 (pS127) in C4-2 cells treated with two independent LATS1/2 siRNAs. (N) Relative growth of C4-2 cells with control or the indicated LATS1/2 siRNA. (O, P) Western blot analysis of LATS1, LATS2, and YAP in (O) or relative growth of (P) C4-2 cells treated with the indicated siRNA. Results in A-H, L, N, and P are representatives of three independent experiments. Data are ± SD. *P<0.05, **P<0.01, ***P<0.001 (Student’s t-test).

Of note, C4-2 was derived from LNCaP in castrated mice and exhibited castration resistance (Wu et al. 1994), whereas 22RV1 and R1-D567 contained AR splicing variants V7 and V567ES, respectively, both of which confer resistance to AR inhibitors such as Enzalutamide (Liu *et al*, 2008; Taylor *et al*, 2010; Ware *et al*, 2014). Consistent with these previous findings, we found that C4-2 was less sensitive to Enzalutamide compared with LNCaP while both 22RV1 and R1-D567 were resistant to Enzalutamide (Fig. 1E-H).

### YAP inhibits AR transcriptional program

AR transcriptional activity is essential for the growth of AR^+^ PCa (Dai *et al*., 2017). To determine whether Hippo pathway inhibition regulates the AR transcriptional program, we examined the transcriptome of C4-2 cells treated with XMU-MP-1 or vehicle by RNA-seq. Gene Ontology (GO) enrichment analysis of differentially expressed genes revealed that AR responsive genes are among the top downregulated transcriptional programs in XMU-MP-1 treated C4-2 cells (Fig. 2A). Gene Set Enrichment Analysis (GSEA) indicated that the AR responsive genes were enriched in genes depleted in XMU-MP-1 treated cells while YAP signature genes were enriched in the genes upregulated by XMU-MP-1 (Fig. 2B-C). RT-qPCR confirmed that XMU-MP-1 treatment inhibited the expression of multiple AR target genes including *KLK2*, *KLK3*, *TMPRSS2*, and *FKBP5* in C4-2, LNCaP, 22RV1 or R1-D567 cells in a dose-dependent manner (Fig. 2D-F; Fig. EV3A-F). XMU-MP-1 also inhibited Dihydrotestosterone (DHT)-induced expression of AR target genes in LNCaP cells similar to Enzalutamide (Fig. 2F). LATS1/2 KD inhibited the expression of AR target genes in C4-2, LNCaP, and 22RV1 cells (Fig. 2H; Fig. EV3H-I). Knockdown of YAP alleviated the inhibitory effect of XMU-MP-1 or LATS1/2 KD on AR target gene expression in C4-2 cells (Fig. 2G, I). On the other hand, overexpression of YAP-WT or YAP-5SA inhibited AR target gene expression in C4-2 and 22RV1 with YAP-5SA exhibiting more potent inhibitory effect than YAP-WT (Fig. 2J-M), consistent with YAP-5SA being more potent in blocking AR^+^ PCa cell growth than YAP-WT (Fig. 1A-D).

**Figure 2.**
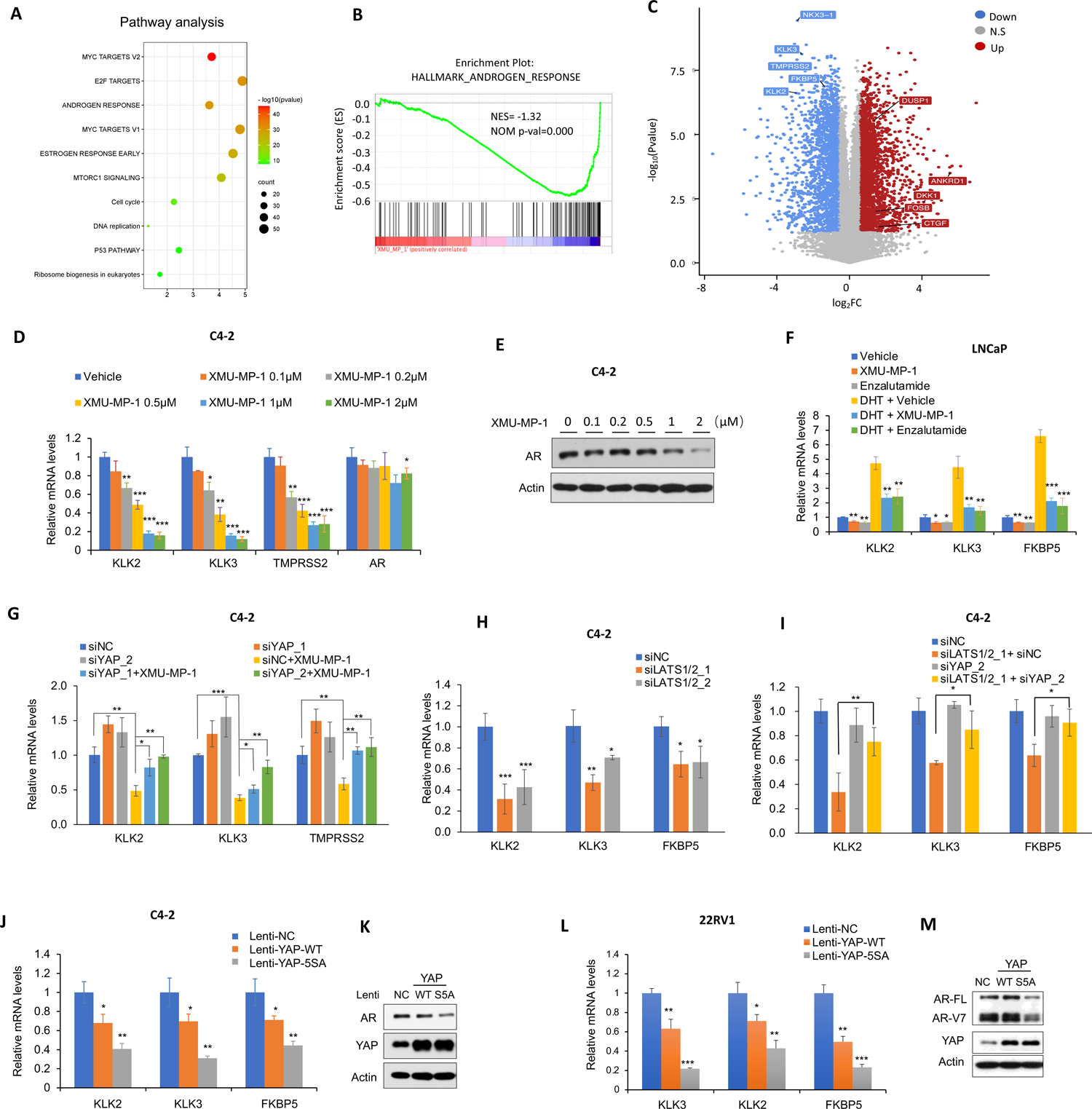
MST1/2 inhibition or YAP activation inhibits AR transcriptional program. (A) Top 10 signaling pathways downregulated in C4-2 cells treated with XMU-MP-1. The cells were treated with vehicle or 2 μM XMU-MP-1 for 8 hours. Threshold P<0.001 and fold change>2. n=3. (B) Gene set enrichment analysis shows a depletion of androgen responsive signature genes in C4-2 cells treated with XMU-MP-1. (C) Volcano plot shows the opposite effects of XMU-MP-1 treatment on androgen responsive signature genes (blue) and the Hippo pathway signature genes (red) in C4-2 cells treated with XMU-MP-1.Threshold P<0.05 and fold change>1.5. (D) Relative mRNA levels of AR and the indicated AR target genes in C4-2 cells treated with increasing concentrations of XMU-MP-1. (E) Western blot analysis of AR in C4-2 cells treated with XMU-MP-1 at the indicated concentrations. (F) Relative mRNA levels of the indicated AR target genes in LNCaP cells grown in the absence or presence 10 nM DHT and treated with vehicle, 2 μM XMU-MP-1, or 20μM Enzalutamide. (G) Relative mRNA levels of the indicated AR target genes in C4-2 cells treated with control (siNC) or YAP siRNA (siYAP_1 or siYAP_2) and with or without 2 μM XMU-MP-1. (H) Relative mRNA levels the indicated AR target genes in C4-2 cells treated with control (siNC) or LATS1/2 siRNA (siLATS1/2_1 or siLATS1/2_2). (I) Relative mRNA levels the indicated AR target genes in C4-2 cells treated with the indicated siRNA. (J-M) Relative mRNA levels of the indicated AR target genes (J, L) or western blot analysis of AR and YAP (K, M) in C4-2 (J, K) or 22RV1 (L, M) cells expressing control (NC), YAP-WT or YAP-5SA constructs through lentiviral infection. Results in D, F-J, and L are representatives of three independent experiments. Data are ± SD. *P<0.05, **P<0.01, ***P<0.001 (Student’s t-test).

### YAP inhibits AR binding to its target promoters

We noticed that treatment of C4-2 and other AR^+^ PCa cells reduced AR protein levels especially at high doses even though AR mRNA levels remained relatively unchanged (Fig. 2D-E; Fig. EV3A-F), suggesting that XMU-MP-1 may regulate AR at a posttranscriptional level. Treating cells with the proteasome inhibitor MG-132 blocked XMU-MP-1-induced downregulation of AR protein levels (Fig. 3A), suggesting that XMU-MP-1 promotes AR protein degradation through the ubiquitin-proteasome pathway. However, blocking AR degradation by MG-132 did not rescue XMU-MP-1-induced downregulation of AR target gene expression (Fig. 3B-C), suggesting that AR protein degradation is not the primary cause of AR transcriptional program downregulation by Hippo pathway inhibition.

**Figure 3.**
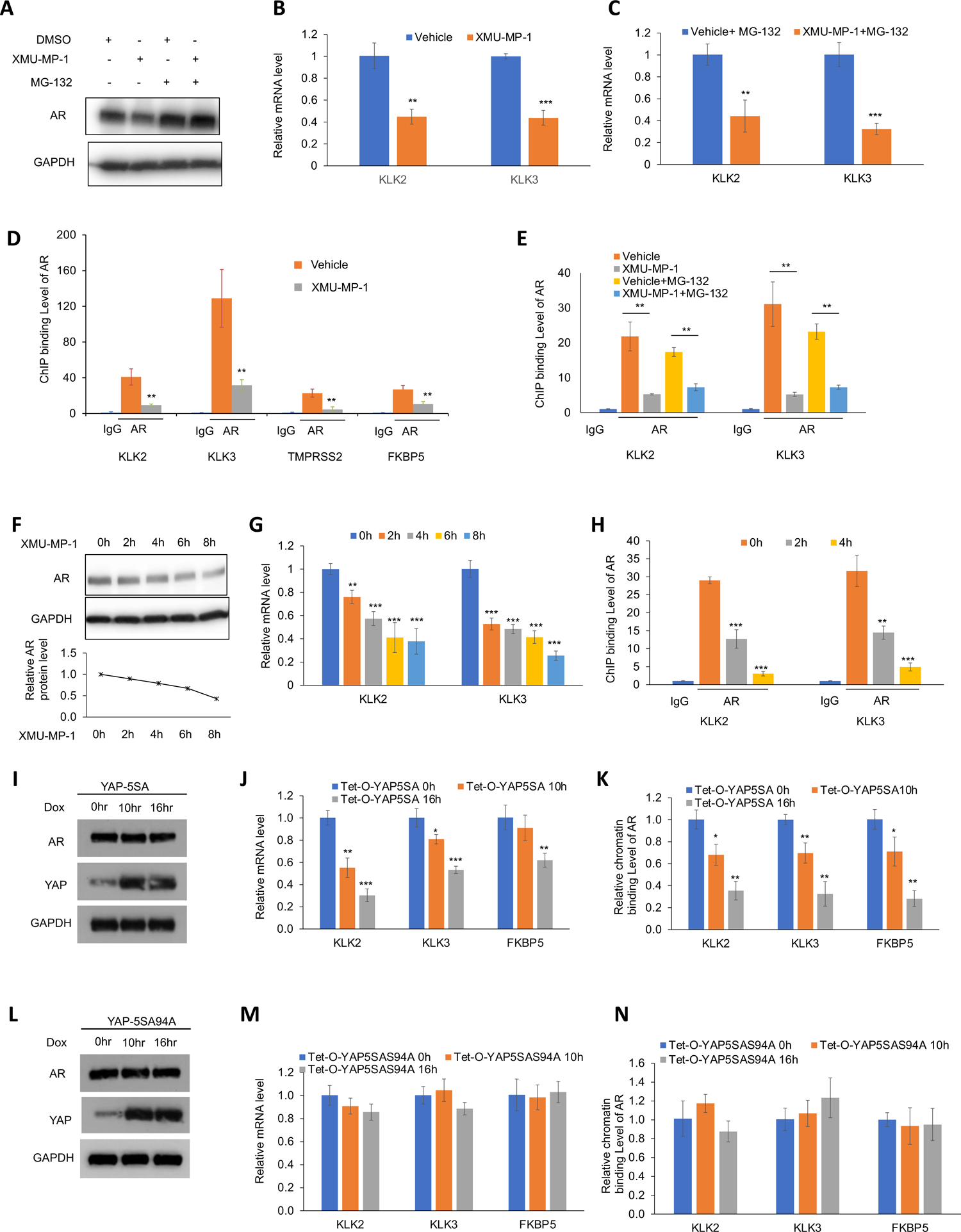
MST1/2 inhibition or YAP overexpression inhibits AR binding to its target promoters/enhancers. (A) Western blot analysis of AR in C4-2 cells treated with DMSO or 2 μM XMU-MP-1 in the absence of presence of 20 nM MG132 for 8 hours. (B-C) Relative mRNA levels of the indicated AR target genes in C4-2 cells treated with DMSO or 2 μM XMU-MP-1 in the absence (B) of presence (C) of 20 nM MG132 for 8 hours. (D) ChIP-qPCR analysis of AR binding to the promoter/enhancer regions of the indicated AR target genes in C4-2 cells treated with vehicle or 2 μM XMU-MP-1 for 4 hours. (E) ChIP-qPCR analysis of AR binding to the promoter/enhancer regions of the indicated AR target genes in C4-2 cells treated with vehicle or 2 μM XMU-MP-1 in the absence of presence of 20 nM MG132 or 8 hours. (F-H) Western blot analysis of AR (F), relative mRNA levels of the indicated AR target genes (G) and ChIP-qPCR analysis of AR binding to the promoter/enhancer regions of the indicated AR target genes (H) in C4-2 cells treated with 2 μM XMU-MP-1 for the indicated periods of time. (I-N) Western blot analysis of AR and YAP (I, L), relative mRNA levels of the indicated AR target genes (J, M) and ChIP-qPCR analysis of AR binding to the promoter/enhancer regions of the indicated AR target genes (K, N) in C4-2 cells stably expressing Tet-O-YAP-5SA (I-K) or Tet-O-YAP-5SAS94A and treated with 0.2 μg/ml doxycycline (Dox) for the indicated periods of time. Results in B-E, G-H, J-K and M-N are representatives of three independent experiments. Data are ± SD. *P<0.05, **P<0.01, ***P<0.001 (Student’s t-test).

To determine the mechanism by which Hippo signaling inhibits AR-mediated transcription, we carried out chromatin immunoprecipitation (ChIP) experiments to determine the binding of AR to the promoter/enhancer regions of its target genes. We found that XMU-MP-1 treatment reduced the occupancy of AR on the promoter/enhancer regions of multiple AR target genes including *KLK2*, *KLK3*, *TMPRSS2* and *FKBP5* (Fig. 3D). Even when AR protein degradation was blocked by MG-132, AR binding to its target promoters was still inhibited by XMU-MP-1 (Fig. 3E). Treating cells with XMU-MP-1 for shorter periods of time (2∼4 hours) also inhibited AR binding to its target promoters as well as AR target gene expression without significantly reducing AR protein levels (Fig. 3F-H). Hence, it appears that inhibition of AR binding to its target promoters proceeded AR protein degradation induced by XMU-MP-1.

To further determine how YAP regulates AR transcriptional activity, we generated C4-2 cell lines that stably expressed an inducible YAP-5SA (Tet-O-YAP-5SA). Dox-induction of YAP-5SA for 10 or 16 hours inhibited AR binding to its target promoters and AR target gene expression with little if any effects on AR protein levels (Fig. 3I-K). Thus, YAP inhibits AR binding to its target promotes, which may subsequently lead to AR degradation. To determine whether YAP inhibits AR by directly binding to it, we carried out CoIP experiments. However, we failed to observe AR association with either YAP-5SA or endogenous YAP (see below). We concluded that YAP inhibits AR by binding to other proteins.

In the canonical Hippo signaling pathway, YAP forms a complex with the pathway transcription factor TEAD to regulate Hippo target gene expression (Wu *et al*., 2003; Zhang *et al*., 2008; Zhao *et al*., 2008). To determine whether YAP inhibits AR depending on its binding to TEAD, we generated C4-2 cell lines that stably expressed a TEAD binding defective form of YAP-5SA (Tet-O-YAP-5SAS94A) (Zhao *et al*., 2008). Dox-induced YAP-5SAS94A expression for 10 or 16 hours neither inhibited AR binding to its target promoters nor affected AR target gene expression (Fig. 3L-N). Hence, YAP inhibits AR transcriptional activity through binding to TEAD.

### TEAD formed a complex with AR to promote AR activity and AR^+^ PCa growth

TEAD family of transcription factors contains four members (TEAD1, TEAD2, TEAD3 and TEAD4) that often function redundantly or additively to regulate Hippo pathway target gene expression (Currey *et al*, 2021; Lin *et al*, 2017). To determine the role of TEAD in AR^+^ PCa, we simultaneously knocked down TEAD1, TEAD3 and TEAD4 using two independent siRNAs (siTEAD1/3/4_1 and siTEAD1/3/4_2) that target shared sequences among these TEAD family members (Zhao *et al*., 2008). TEAD1/3/4 knockdown inhibited the expression of AR target gene in C4-2, 22RV1 and R-D567 cells as well as the growth of these AR^+^ PCa cells (Fig. 4A-C; Fig. EV4A-F). On the other hand, TEAD4 or TEAD1 overexpression increased the AR target gene expression and alleviated the inhibition of AR target gene expression by XMU-MP-1 in C4-2 cells (Fig. 4D; Fig. EV4G), suggesting that TEAD positively regulates AR-mediated transcription, which is opposite to YAP.

**Figure 4.**
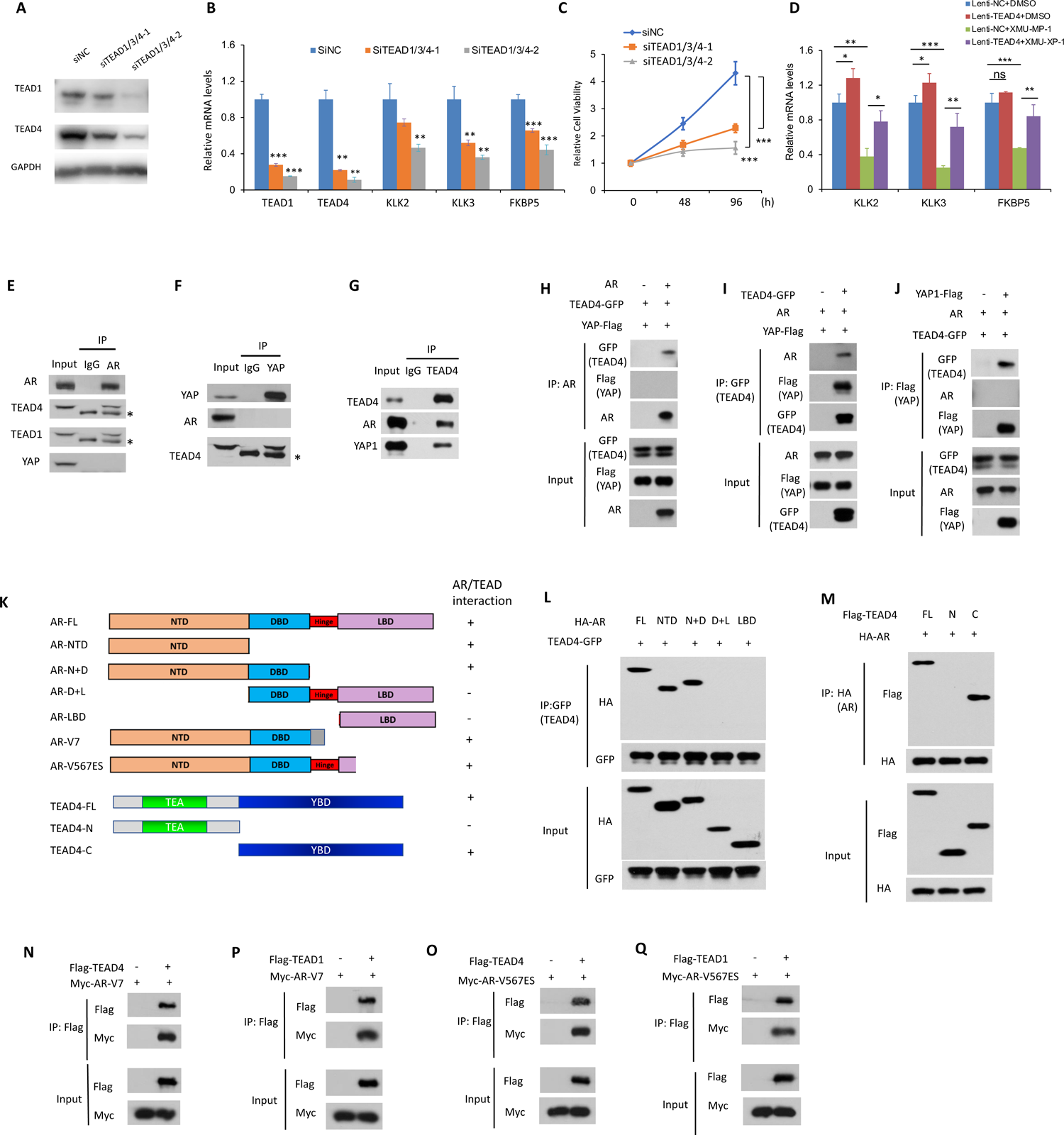
TEAD forms a complex with AR and promotes AR signaling. (A-C) Western blot analysis of TEAD1 and TEAD4 (A), relative mRNA levels of TEAD1 and TEAD4 and the indicated AR target genes (B) and relative growth of C4-2 cells treated with two independent TEAD1/3/4 siRNA or control siRNA. (D) Relative mRNA levels of the indicated AR target genes in C4-2 cells expression the control or TEAD4 constructs and treated with vehicle or XUM-MP-1. (E-G) C4-2 cell extracts were immunoprecipitated with IgG, anti-AR (E), anti-YAP (F) or anti-TEAD4 (G) antibodies. Immunoprecipitates and cell lysates (input) were analyzed by western blot with the indicated antibodies. Asterisks in E and F indicate IgG. (H-J) HEK-293T cells transfected with the indicated constructs. Cell lysates were immunoprecipitated with the indicate antibodies. Immunoprecipitates and cell lysates (input) were analyzed by western blot with the indicated antibodies. (K) Schematic diagrams of the indicated AR and TEAD constructs. (L-Q) HEK-293T cells transfected with the indicated AR, TEAD and YAP constructs. Cell lysates were immunoprecipitated with the indicate antibodies. Immunoprecipitates and cell lysates (input) were analyzed by western blot with the indicated antibodies. Results in in B-J and L-Q are representatives of three independent experiments. Data are ± SD. *P<0.05, **P<0.01, ***P<0.001 (Student’s t-test).

CoIP experiments revealed that endogenous AR interacted with both TEAD1 and TEAD4 but not with YAP in C4-2 cells whereas endogenous YAP formed a complex with TEAD1 and TEAD4 but not with AR in C4-2 cells (Fig. 4E-F). As expected, endogenous TEAD4 formed a complex with AR or YAP in C4-2 cells (Fig. 4G). Similar results were obtained when these proteins were coexpressed in HEK-293T cells (Fig. 4H-J). These results demonstrate that AR/TEAD and YAP/TEAD form distinct protein complexes.

Deletion mapping revealed that the C-terminal half of TEAD (YBD: YAP binding domain) and the N-terminal domain (NTD) of AR mediated the association between TEAD and AR (Fig. 4K-M). 22RV1 and R-D567 cells express the AR splicing variants AR_V7 and AR_V567ES, respectively, both of which have their ligand binding domain (LBD) deleted but contain the intact NTD (Fig. 4K). Consistent with AR binding to TEAD through its NTD, we found that both AR_V7 and AR_V567ES formed a complex with TEAD4 or TEAD1 when coexpressed in HEK-293T cells (Fig. 4N-Q). These results suggest that TEAD influences AR activity and AR^+^ PCa growth by physically interacts with AR.

### TEAD promotes AR binding to its target promoters/enhancers

Next, we examined whether TEAD and AR form a complex on chromatin by carrying out ChIP-seq experiments. We found that AR and TEAD co-bound 14857 peaks on chromatin, which represent 24% of the total AR binding peaks (62304) and 55% of total TEAD binding peaks (26807) and include those in the promoter/enhancer regions of *KLK2*, *KLK3*, and *FKBP5* (Fig. 5A-D). Motif analysis indicates that both TEAD binding sites and AR binding sites (ARE) are enriched in the AR and TEAD co-binding (AR+/TEAD+) peaks whereas AR binding alone (AR+/TEAD-) and TEAD binding alone (AR-/TEAD+) peaks only contain ARE or TEAD binding motifs, respectively (Fig 5E). All three classes of binding peaks are enriched in the binding motifs for FOXA1, FOXA2, and FOXM1 (Fig. 5E), consistent with previous findings that these transcription factors act as pioneer factors to regulate the chromatin accessibility of other transcription factors (Friedman & Kaestner, 2006). Combined analysis of RNA-seq and ChIP-seq data revealed that, among 100 well annotated AR target genes (Liberzon *et al*, 2015), 42 were downregulated by XMU-MP-1 in C4-2 cells (Table EV1), and that >90% (38/42) of these AR target genes contain AR and TEAD co-binding peaks in their promoter/enhancer regions (Fig. 5D, F; Table EV2). In addition, ∼40% (848/2136) of genes downregulated by XMU-MP-1 in C4-2 cells contain AR and TEAD co-binding peaks in their promoter/enhancer regions (Fig. 5G). ChIP-qPCR experiments validated the co-binding of AR and TEAD4 on the promoter/enhancer regions of AR target genes including *KLK2*, *KLK3*, and *FKBP5* in C4-2 cells (Fig. 5H). Knockdown of TEAD1/3/4 reduced the occupancy of AR on these target promoters (Fig. 5I), and reciprocally, TEAD4 binding to the promoter/enhancer regions of AR target genes was also downregulated when AR was knocked down (Fig. 5J). The mutual dependency between AR and TEAD with respect to their optimal promoter occupancy suggest that AR and TEAD bind cooperatively to these target sites.

**Figure 5.**
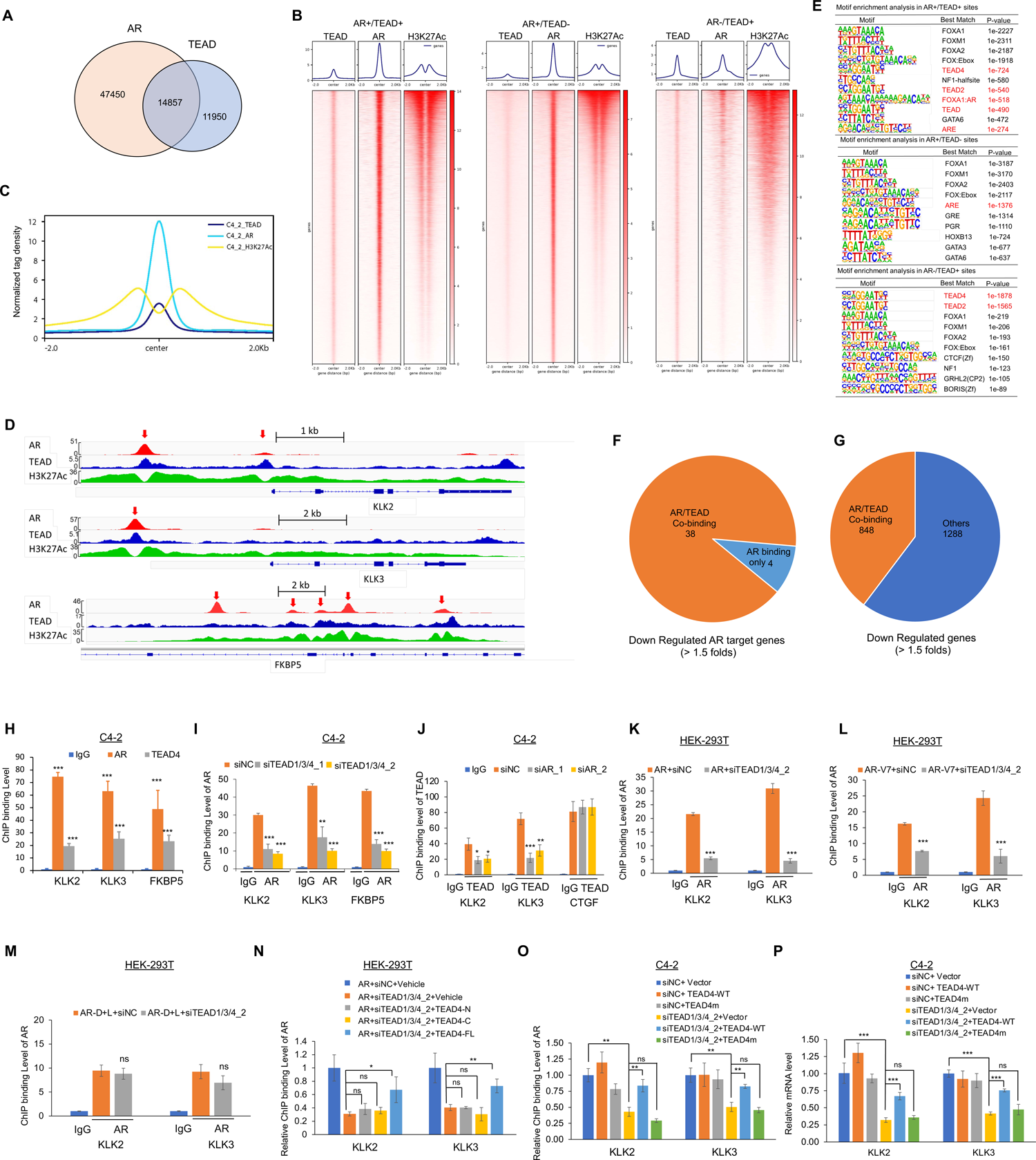
TEAD promotes AR binding to its target promoters/enhancers. (A) Venn diagram showing the numbers of AR binding peaks overlapping with TEAD binding peaks in C4-2 cells in ChIP-Seq experiments. (B) Heatmaps of TEAD, AR, and H3K27Ac ChIP-Seq reads from C4-2 cells showing the bound loci of AR^+^/TEAD^+^, AR^+^/TEAD^-^ and AR^-^/TEAD^+^. (C) Aggregate plots showing the normalized tag density of TEAD, AR, and H3K27Ac at indicated gene binding sites within 2kb frames in C4-2 cells. (D) ChIP-seq signal tracks of AR, TEAD, and H3K27Ac at promoter/enhancer regions of the indicated AR target genes in C4-2 cells. (E) Transcription factor binding motif enrichment analyses of AR^+^/TEAD^+^, AR^+^/TEAD^-^, and AR^-^/TEAD^+^ sites. (F) >90% (38/42) of AR target genes downregulated by XMU-MP-1 in C4-2 cells are co-bound by both TEAD and AR in their promoter/enhancer regions. (G) ∼40% (848/2136) of genes by XMU-MP-1 in C4-2 cells are co-bound by both TEAD and AR. (H) ChIP-qPCR analysis shows that TEAD4 co-binds with AR to the promoter/enhancer regions of the indicated AR target genes in C4-2 cells. (I) ChIP-qPCR analysis of AR binding to the promoter/enhancer regions of the indicated AR target genes in C4-2 cells treated with control (siNC) or two independent TEAD1/3/4 siRNAs. (J) ChIP-qPCR analysis of TEAD binding to the indicated AR targets in C4-2 cells treated with control (siNC) or two independent TEAD1/3/4 siRNAs. (K-M) ChIP-qPCR analysis of *KLK2* and *KLK3* promoter/enhancer occupancy of full-length AR (K), AR-V7 (L) or AR-D+L (M) exogenously expressed HEK-293T cells treated with control or TEAD siRNA. (N) ChIP-qPCR analysis shows that exogenous expression of full-length TEAD4 (TEAD4-FL) but not truncated TEAD4 (TEAD4-N and TEAD4-C) partially rescued of AR binding to *KLK2* and *KLK3* promoters in HEK-293T cells with endogenous TEAD depleted by siTEAD1/3/4_2. (O-P) ChIP-qPCR (O) and RT-PCR analyses show that exogenous expression of wild type (TEAD4-WT) but not the DNA binding deficient form (TEAD4m) of TEAD4 rescued AR binding to the promoter/enhancer regions of *KLK2* and *KLK3* (O) as well as the expression of these genes (P) in C4-2 cells treated with TEAD1/3/4 siRNA. Results in H-P are representatives of three independent experiments. Data are ± SD. *P<0.05, **P<0.01, ***P<0.001 (Student’s t-test).

To further determine whether TEAD-AR interaction and TEAD DNA binding are important for AR to bind its target sites, we turned to HEK-293T cells. We found that knockdown of TEAD1/3/4 reduced the binding of exogenously expressed full-length AR (AR-FL) and AR splicing variant AR-V7 to the promoter/enhancer regions of *KLK2* and *KLK3* (Fig. 5K-L). By contrast, siTEAD1/3/4 did not affect the promoter occupancy of the exogenously expressed AR deletion mutant (AR_D+L) that no longer interacted with TEAD (Fig. 5M). Exogenous expression of an RNAi insensitive full-length TEAD4 rescued AR-FL binding to its target sites in TEAD1/3/4-depleted cells; however, expression of RNAi insensitive TEAD4 mutants lacking either the AR binding domain (TEAD4-N) or the DNA binding domain (TEAD4-C) failed to rescue AR binding to its target sites (Fig. 5N). We also generated a DNA binding deficient full-length form of TEAD4 (TEAD4m) in which critical residues that contact both the major and minor groves of DNA were mutated: R64/K65/I66/L68A/ R95/K96/S100/Q103A (Shi *et al*, 2017). Unlike the wild type TEAD4 (TEAD4-WT), the DNA binding deficient TEAD4m nether rescued AR binding to its target promoters/enhancers nor AR target gene expression in TEAD1/3/4 depleted C4-2 cells (Fig. 5O-P). Taken together, these results suggest that both TEAD/AR interaction and TEAD DNA binding are required for optimal binding of AR to its target promoters/enhancers.

### YAP inhibits AR-TEAD complex formation by competing for TEAD

The observation that both YAP and AR bind to the same region in TEAD raised a possibility that YAP binding to TEAD may prevent AR from binding to TEAD. Indeed, coexpression of increasing amounts of YAP with fixed amounts of AR and TEAD4 in HEK-293T cells progressively reduced the amounts of AR associated with TEAD4 with concomitant formation of increasing amounts of YAP-TEAD4 complex (Fig. 6A). Treating C4-2 cells with XMU-MP-1 increased the amount of YAP-TEAD4 complex while decreased the amount of AR associated with TEAD4 (Fig. 6B). Dox-induced YAP-5SA but not YAP-5SAS94A inhibited the formation of AR-TEAD4 complex formation in C4-2 cells (Fig. 6C), which correlates with their ability or inability to inhibit AR binding to its target promoters and AR target gene expression (Fig. 3I-N). Furthermore, Dox-induction of YAP-5SA in C4-2 cells inhibited co-binding of AR and TEAD on AR target promoters whereas TEAD binding to the canonical YAP target gene CTGF was enhanced (Fig. 6D, E). When coexpressed in HEK-293T cells, YAP-5SA inhibited the promoter/enhancer occupancy of AR-FL and AR-V7 but not of AR_D+L on AR target genes *KLK2* and *KLK3* (Fig. 6F-H), and that coexpression of TEAD4 partially rescued the promoter/enhancer occupancy of AR-FL inhibited by YAP-5SA (Fig. 6I). Taken together, these results suggest that YAP and AR compete for TEAD and that YAP binding to TEAD interfere with TEAD/AR interaction and their cooperative binding to AR target promoters, leading to diminished AR transcriptional activity (Fig. 6J).

**Figure 6.**
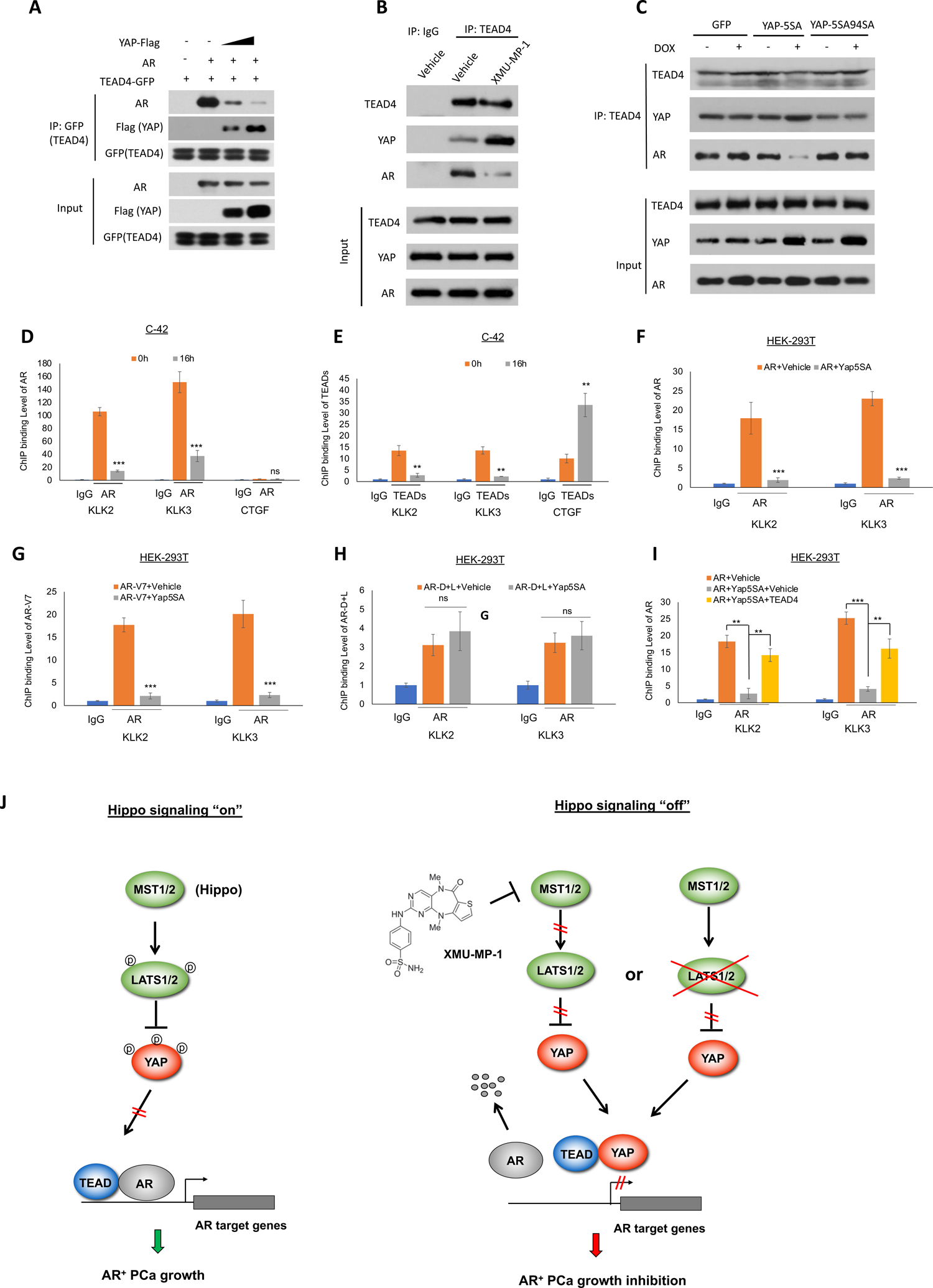
YAP binding to TEAD impedes AR/TEAD cooperativity. (A) CoIP experiments with the indicated antibodies followed by western blot using cell extracts from HEK-293T cells transfected with the indicated constructs show that increasing amounts of YAP displaced AR from TEAD4. (B) XMU-MP-1 increased YAP but decreased AR binding to TEAD4. C4-2 cell were treated with vehicle or 2 μM XMU-MP-1 for 4 hours. Cell extracts were immunoprecipitated with IgG or anti-TEAD4 antibody. Immunoprecipitates and cell lysates (input) were analyzed by western blot with the indicated antibodies. (C) YAP-5SA but not YAP-5SAS94A inhibited AR/TEAD interaction. C4-2 cells stably expressing Tet-O-GFP, Tet-O-YAP-5SA or Tet-O-YAP-5SAS94A were treated with 0.2 μg/ml doxycycline (Dox) for 10 hours, followed by CoIP and western blot analyses with the indicated antibodies. (D-E) ChIP-qPCR analysis of AR (D) and TEAD (E) binding to the promoter/enhancer regions of *KLK2*, *KLK3* and *CTGF* in Tet-O-YAP-5SA expressing C4-2 cells treated with 0.2 μg/ml doxycycline (Dox) for the indicated time. (F-H) ChIP-qPCR analysis of AR binding to the promoter/enhancer regions of *KLK2* and *KLK3* in HEK293 cells transfected with YAP-5SA and the indicated AR constructs. (I) ChIP-qPCR analysis to show that coexpression of TEAD4 attenuated the YAP-5SA-mediated inhibition of AR binding to the promoter/enhancer regions of *KLK2* and *KLK3* in HEK293 cells. (J) Working model for Hippo and AR signaling crosstalk. MST1/2 inhibits YAP nuclear localization through LATS1/2, allowing TEAD to cooperate with AR to drive a transcription program essential for AR^+^ prostate cancer growth. Pharmacological inhibition of MST1/2 promotes YAP nuclear localization and YAP/TEAD interaction, which disrupts TEAD and AR cooperativity, leading to downregulation of AR target gene expression and reduced AR^+^ prostate cancer growth. Results are representatives of three independent experiments Data ± SD. ns: not significant, *P<0.05, **P<0.01, ***P<0.001 (Student’s t-test).

### YAP inhibits hormone therapy resistant AR variants *in vivo*

Changes in AR including amplification, point mutations, and AR variants (AR-Vs) lacking ligand binding domain have been implicated in anti-hormone therapy resistance (Beltran *et al*, 2013; Grasso *et al*, 2012; Henzler *et al*, 2016; Liu *et al*., 2008; Robinson *et al*, 2015; Taylor *et al*., 2010; Ware *et al*., 2014; Watson *et al*, 2015). AR-Vs are thought to be constitutively active and resistant to anti-AR inhibitors such as Enzalutamide due to their loss of the C-terminal LBD (Antonarakis *et al*, 2014; Guo *et al*, 2009; Hu *et al*, 2009; Sun *et al*, 2010). The most common AR-V is AR-V7, which contains exons 1, 2, 3, and a cryptic exon 3b, and the encoded protein lacks the LBD (Zhang *et al*, 2011). Our *in vitro* study showed that MST1/2 inhibition or YAP activation could reduce the growth of CRPC cells (C4-2) or AR splicing variant expressing cells (22Rv1 and R1-D557) while Enzalutamide failed to inhibit the growth of 22RV1 and R1-D557 (Fig. 1E-H). To determine whether YAP activation can inhibit the growth of these anti-hormone therapy resistant PCa cells *in vivo*, we generated xenograft models carrying C4-2 cells that expressed Tet-O-EGFP (control), Tet-O-YAP-5SA, or Tet-O-YAP-5SAS94A, and 22Rv1 cells that expressed Tet-O-YAP-5SA. After tumors grew to 100 mm^3^, mice were treated with either Dox or vehicle (PBS) daily for 3 weeks. We found that Dox-induced YAP-5SA near-completely blocked C4-2 and 22RV1 tumor growth in xenografts whereas the S94 mutation greatly diminished the ability of YAP to inhibit C4-2 tumor growth (Fig. 7A-F; Fig. EV5A-C), suggesting that YAP can potently inhibit AR^+^ prostate cancer growth *in vivo* in a manner depending on its binding to TEAD. Of note, YAP-5SAS94A still slightly inhibited C4-2 tumor growth in xenografts (Fig. EV5A-C), likely because S94A mutation did not completely abolish YAP-TEAD interaction and prolonged expression of YAP-5SAS94A might still be able to inhibit AR signaling (Li *et al*, 2010). Indeed, examination of the tumors expressing YAP-5SAS94A for 3 weeks showed that AR target genes were inhibited, albeit less dramatically compared with YAP-5SA expressing tumors (Fig. EV5D).

**Figure 7.**
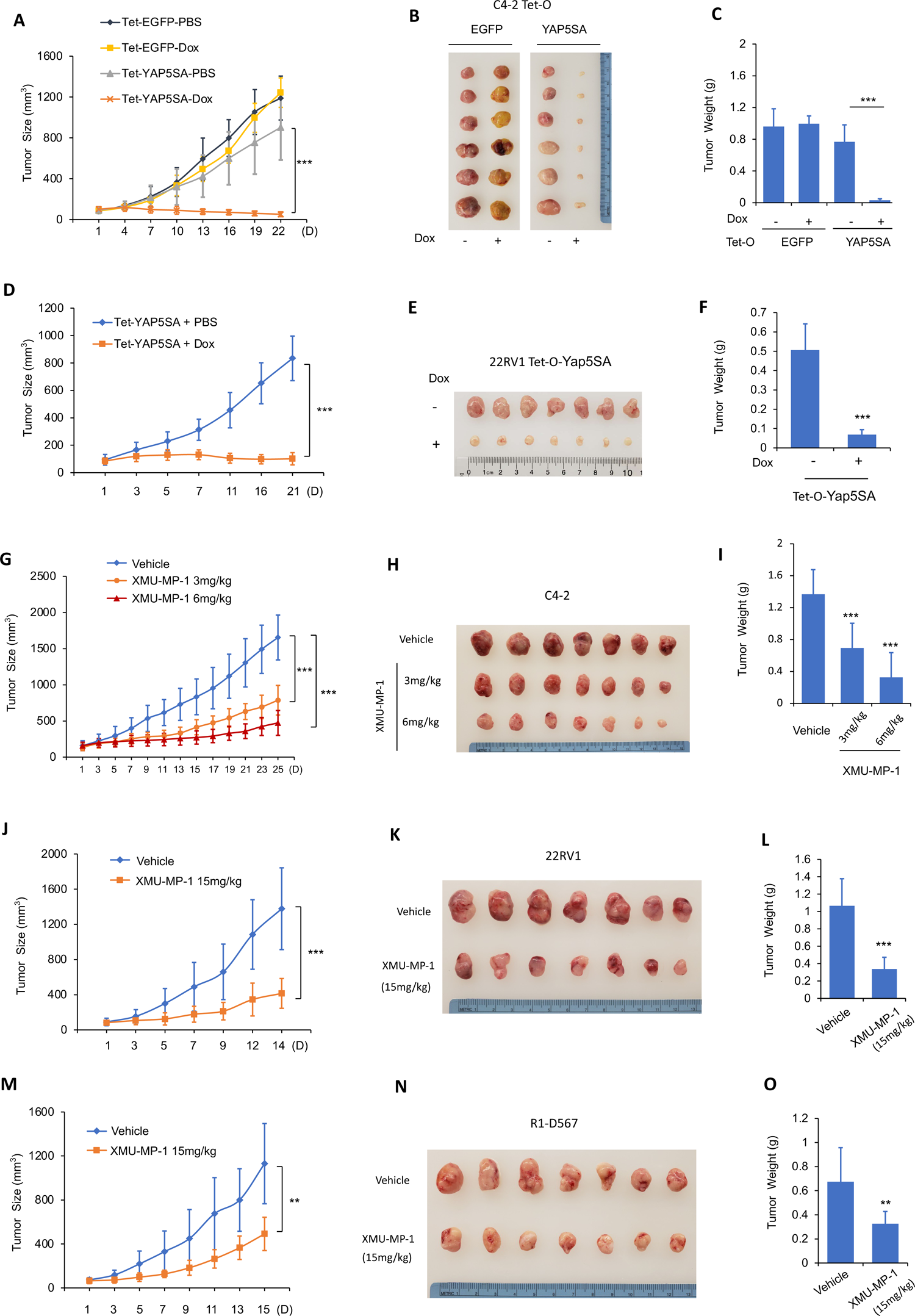
MST1/2 inhibition/YAP activation inhibits therapy resistant AR *in vivo*. (A-F) YAP activation inhibited AR positive prostate cancer growth in xenografts bearing C4-2 or 22RV1 tumors that stably express Tet-O-YAP-5SA. Male NOD scid gamma (NSG) mice bearing C4-2 tumor cells stably expressing a Tet-O-EGFP construct were used as control. Male NOD scid gamma (NSG) mice bearing the indicated tumors were injected i.p. with PBS containing Dox (20mg/kg) or PBS daily for the indicated period of time. Tumor growth curve (A, D), photograph of tumor samples (B, E), and quantification of tumor weight (C, F) at the end of treatment were shown. (G-O) XMU-MP-1 inhibited AR positive prostate cancer growth in xenografts bearing C4-2 (G-J), 22RV1 (J-L) or R1-D567 (M-O) tumors. Male NOD scid gamma (NSG) mice bearing the indicated tumors were treated daily with XUM-XP-1 at the indicated concentrations or with vehicle for the indicated periods of time. Tumor growth curve (G, J, M), images of tumor samples (H, K, N), and quantification of tumor weight (I, L, O) at the end of treatment were shown. Data are ± SD. **P<0.01, ***P<0.001 (Student’s t-test). n=7 mice for each group.

We next determined whether pharmacological inhibition of MST1/2 by XMU-MP-1 could inhibit the growth of therapy resistant PCs *in vivo*. Xenografts carrying C4-2 cells were treated with two different doses of XUM-MP-1 (3mg/kg or 6mg/kg daily) or vehicle for 25 days. As shown in Fig. 7G-I, XUM-MP-1 dramatically reduced the tumor growth in a dose-dependent manner. We also found that XUM-MP-1 could inhibit tumor growth in xenografts expressing AR variants V7 (22RV1) and V567ES (R1-D567) albeit less effectively compared with its ability to inhibit C4-2 tumor growth (Fig. 7J-O). Consistent with the previous study that XUM-MP-1 was well tolerated in mice (Fan *et al*., 2016), we found that mice treated with XUM-MP-1 exhibited normal body weight and liver size except that spleens were slightly increased when treated with high doses of XUM-MP-1 (Fig. EV6). We also treated 22RV1-tumor bearing mice with XMU-MP-1 or Enzalutamide side by side and found that Enzalutamide had little if any effect on 22RV1 tumor growth whereas XMU-MP-1 could dramatically inhibit the tumor growth (Fig. EV7).

## Discussion

Although Hippo signaling pathway mostly functions as a tumor suppressor pathway and YAP as an oncoprotein to drive cancer progression and confer cancer drug resistance, recent studies suggest YAP could function as a tumor suppressor in several contexts (Cheung *et al*., 2020; Cottini *et al*., 2014; Li *et al*., 2020; Ma *et al*., 2021). Here we demonstrated that YAP activation by pharmacological inhibition of Hippo/MST1/2 kinase activity, LATS1/2 KD, or transgenic expression of YAP-5SA inhibited AR transcriptional activity and AR^+^ PCa growth. By exploring the underlying mechanism, we uncovered a noncanonical mechanism by which YAP acts through TEAD to regulate cancer cell growth. In the conventional model, YAP binding to TEAD either activates oncogenic programs or inhibits the transcription of tumor suppressors (Kim *et al*, 2015; Zanconato *et al*, 2015). Here, we showed that TEAD is a critical cofactor for AR, which is the major oncogenic driver in AR^+^ PCa. TEAD and AR co-bind a large set of AR target promoters/enhancers to promote AR transcriptional activity. YAP and AR compete for TEAD and YAP binding to TEAD disrupts the cooperativity between TEAD and AR, leading to AR dissociation from its target promoters/enhancers and subsequently proteasome-mediated degradation, diminished AR target gene expression, and inhibition of AR^+^ PCa growth (Fig. 6J). Hence, instead of acting in the same direction described by most of the literatures (Currey *et al*., 2021; Lin *et al*., 2017), YAP and TEAD have an antagonistic relationship in the regulation of AR transcriptional activity and AR^+^ PCa growth. Interestingly, our recent study has shown that TEAD can also function as a cofactor for estrogen receptor (ER) and that YAP inhibits ER^+^ breast cancer by blocking TEAD/ER interaction (Li *et al*, 2022). Therefore, our findings that YAP inhibits cancer growth by interfering with the cooperativity between TEAD and an oncogenic driver may represent a general mechanism that can be applied to multiple cancers.

A previous study suggested that YAP forms a complex with AR to promote AR signaling and prostate cancer growth (Kuser-Abali *et al*, 2015). However, we did not detect any interaction between YAP and AR expressed either endogenously in C4-2 cells or exogenously in HEK-293T cells by CoIP experiments. Instead, we found that AR forms a complex with TEAD, and that YAP competes with AR for binding to the same pool of TEAD. Domain mapping revealed that TEAD binds AR through its C-terminal domain that has been previously shown to bind YAP, which could explain why AR and YAP bindings to TEAD are mutually exclusive. On the other hand, AR interacts with TEAD through its NTD domain that is present in AR splicing variants including AR-V7 and AR-V567ES, which may explain why YAP activation or Hippo pathway inhibition could impede the growth of PCa cells expressing these AR variants. Indeed, we found that both AR-V7 and AR-V567ES interacted with TEAD and that the binding of AR-V7 to AR target promoters/enhancers was downregulated by TEAD knockdown or YAP overexpression.

Prostate cancer is the second most diagnosed malignancy in USA where ∼30,000 men died of metastatic prostate cancer each year (Siegel *et al*., 2018). Most of the prostate cancer is driven by androgen receptor, making the androgen deprivation therapy via pharmacological or surgical castration a stand care of prostate patients. However, a significant portion of treated patients will eventually develop castration-resistant prostate cancer (CRPC). Although the second-generation antiandrogen drugs such as AR antagonist Enzalutamide can extend the survival of metastatic CRPC patients by 4-5 months, primary or acquired resistance to these agents is common among treated patients (de Bono *et al*, 2011; Linder *et al*, 2018; Tran *et al*, 2009). The common mechanisms of Enzalutamide resistance include AR amplification and upregulation of AR variants that lack ligand binding domain such as AR-V7 (Prekovic *et al*, 2018; Wang *et al*, 2021). Our finding that YAP can inhibit AR^+^ prostate cancer cell proliferation by blocking AR transcriptional activity opens an opportunity for developing novel therapeutics to treat AR^+^ prostate cancer patients. Furthermore, our observation that pharmacological inhibition of Hippo/MST1/2 could block the growth of prostate cancer cells that express AR splicing variants suggests that targeting the Hippo pathway may provide a potential therapeutical approach to overcome antiandrogen therapy resistance caused by these variants.

## Materials and Methods

### DNA constructs

The YAP-WT and YAP-5SA constructs were described previously(Cho *et al*, 2018). YAP-5SAS94A was generated by PCR-based mutagenesis from YAP-5SA. For the lentivirus infection, the YAP-WT, YAP-5SA, TEAD4, TEAD4m and TEAD1 were cloned into the pLVX-IRES-ZSgreen vector. To construct Tet-O-EGFP, Tet-O-YAP-5SA and Tet-O-YAP-5SAS94A, GFP, YAP-5SA and YAP-5SAS94A coding sequences were subclone into pTet-O-Ngn2-puro (Addgene plasmid, Cat. No 52047), in which T2A sequence was replaced by UBC promotor. pLenti CMV rtTA3 Hygro (Addgene plasmid, Cat. No. 26730) was used to generate Dox-inducible cell lines. The human TEAD4 constructs were purchased from Origene (https://www.origene.com/; RC219686). TEAD4 full length and deletion constructs were amplified by PCR and the PCR products were sub-cloned in to a pcDNA3.1-FLAG vector. The TEAD1 construct was amplified by PCR and sub-cloned in to a p3xflag-CMV-10 vector. AR-WT construct was acquired from Addgene (pCMV-hAR, Addgene plasmid, Cat. No 89078). The AR-FL and deletion mutants and spicing variants constructs were amplified by PCR and the PCR products were sub-cloned into pCMV-HA or pCMV-myc vector.

### Cell culture

LNCaP, C4-2, 22RV1 and HEK-293T cells were originally acquired form American Type Culture Collection (ATCC). R1-D567 was generated by a previous study (Nyquist *et al*., 2013; Sramkoski *et al*., 1999). LNCaP, C4-2, 22RV1 and R1-D567 cells were maintained with RPMI-1640 (Gibco, Cat. No. 42401018) supplemented with 2 mM L-glutamine (Gibco, Cat. No. 25030081) and 10% FBS. HEK-293T were cultured with Dulbecco’s Modified Eagle’s Medium that contains 4,5 g/L glucose and 4 mM L-glutamine (DMEM, Gibco, Cat. No. 11965092) supplemented with 10% Fetal Bovine Serum (FBS, Gibco, Cat. No. 16000044). For packaging lentivirus, the HEK-293T cells were transfected with the expression vectors and package vectors (psPAX2 and pMD2.G) by PolyJet (SignaGen laboratories, Cat. No. SL100688). After 48 hours, the viruses were harvested for cell line infection with standard method.

### Reagents

The reagents used were: XMU-MP-1(MCE, Cat. No. HY-100526), Enzalutamide (MCE, Cat. No. HY-70002), 5α-DHT-D_3_ (DHT; Sigma Cat: D-077), Doxycycline (Dox) (Sigma, Cat. No. 33429), Hygromycin B (Sigma, Cat. No. H3274), Puromycin (Sigma, Cat. No. P9620), MG-132 (Calbiochem, Cat. No. 474790).

### Immunoblot analysis

Cells were harvested and lysed with lysis buffer containing 1M Tris pH8.0, 5M NaCl, 1M NaF, 0.1M Na_3_VO_4_, 1% NP-40, 10% glycerol, and 0.5M EDTA (pH 8.0). Proteins were separated by electrophoresis on SDS-polyacrylamide gel electrophoresis (PAGE) and electro-transferred to PVDF membrane. Membranes were washed with TBST and incubated with primary antibodies for 2 hours. And then the membranes were washed for three times with TBST and incubated with second antibodies for 2 hours, after washed for three times with TBST, the membranes were probed with ECL system (Cytiva, Cat. No. RPN2105). The antibodies used in this study were listed here: Anti-YAP (Santa Cruz, Cat. No. sc-101199); Anti-AR (Sigma, Cat. No. A9853); Anti-TEAD4 (Santa Cruz, Cat. No. sc-101184); Anti-TEAD1(BD Transduction Laboratories, Cat. No. 610922); Anti-HA (COVANCE, Cat. No. MMS-101R); Anti-Myc (Abcam, Cat. No. ab32); Anti-Myc (Abcam, Cat. No. Ab9106); Anti-Flag (Sigma, Cat. No. F3165); Anti-GFP (Abcam, Cat. No. ab290); Anti-LATS1 (Cell Signaling Technology, Cat.No. 3477S); Anti-LATS2 (Cell Signaling Technology, Cat.No. 5888S); Anti-YAP (pS109) (Cell Signaling Technology, Cat.No. 53749S); Anti-YAP(pS127) (Cell Signaling Technology, Cat.No.13008S). Peroxidase-Conjugated AffiniPure Goat Anti-Mouse IgG (Jackson ImmunoResearch Code,115-035-003) or Goat Anti-Rabbit IgG (Jackson ImmunoResearch, Code,111-035-144). Chemiluminescent signals were visualized with ECL system (Cytiva, Cat. No. RPN2105).

### Co-immunoprecipitation assay

Immunoprecipitation was performed according to standard protocol. C4-2 cell lysates were incubated with antibodies for AR (Sigma, Cat. No.A9853), YAP (Santa Cruz, Cat. No. SC-101199), TEAD1(BD Transduction Laboratories, Cat. No. 610922) or TEAD4 (Santa Cruz, Cat. No. sc-101184), or mouse IgG (used as the negative control) for overnight while, followed by immobilization and precipitation with Protein A resin. The bound proteins were analyzed by western blot. For overexpression experiments, HEK-293T cells were transfected with 5μg epitope-tagged AR (full length or deletion mutants) and TEAD4 plasmids (full length or deletion mutants) in the absence or presence of Flag-YAP in 10 cm dishes. Cell lysates were incubated with antibodies against epitope tags for overnight, followed by immobilization and precipitation with Protein A resin. The bound proteins were analyzed by immunoblot assay.

### Immunofluorescence assay

C4-2 cells were fixed with 4% paraformaldehyde in PBS for 10 min, permeabilized with 0.2% Triton X-100 (Sigma, Cat. No. T8787) for 5 min and blocked by 5% BSA in PBS for 1 h. A rabbit anti-AR antibody (Cell signaling, Cat. No. 5153) and mouse anti-YAP monoclonal antibodies (Santa Cruz, Cat. No. SC-101199) were used, followed by Cy2- and Cy3-conjugated secondary antibodies (Jackson ImmunoResearch). As negative controls, the samples were incubated with the secondary antibodies without primary antibodies. Images were captured by Zeiss LSM510 confocal microscopy. The acquired pictures were further processed and assembled using ImageJ.

### RNA interference

For RNAi experiments in prostate cancer cells, the siRNAs were acquired from the Sigma-Aldrich. The sequences for YAP silencing were: 5’-CAC CUA UCA CUC UCG AGA U-3’ and 5’-GCU CAU UCC UCU CCA GCU U-3’. The sequences for TEAD1/3/4 silencing were: 5’-AUG AUC AAC UUCA UCC ACA AG-3’ and 5’ GAU CAA CUU CAU CCA CAA GCU-3’. The sequences for AR silencing were: 5’-CCA UCU UUC UGA AUG UCC U-3’ and 5’-CAG GAA UUC CUG UGC AUG A-3’. The sequences for negative control were: 5’-UUC UCC GAA CGU GUC ACG U-3’. The LATS1/2 siRNAs were purchased from Sigma, the siRNA ID of siLATS1 were SASI_Hs01_00046128 and SASI_Hs01_00046129, the siRNA ID of siLATS2 were SASI_Hs01_00158803 and SASI_Hs01_00158806. The RNAi MAX reagent (Invitrogen Cat. No. 13778150) was used for the transfection of siRNA. The knockdown efficiency was validated via real-time PCR and immuno-blotting.

### Virus infection and transient transfection

For the virus infection, the expression vectors were transfected together with package vectors (psPAX2 and pMD2.G) into HEK-293T cells by PolyJet (SignaGen laboratories, Cat. No. SL100688). After 48 hours, the supernatants of the medium were collected and filtered with 0.45 μm filter. The supernatant containing virus was stored in 4℃ for cell infection. The prostate cancer cells were cultured in fresh media and subsequently infected with lentivirus overnight together with Polybrene (Sigma, Cat. No. H9268). For cell transfection, PolyJet was utilized according to the protocol.

### RNA extraction and RT-qPCR analysis

We extracted the total RNA by RNeasy plus mini kits according to the protocol (Qiagen, Cat. No. 74106). After RNA extraction, the RNA was subjected to reverse transcription PCR for cDNA synthesis according to the RT-PCR kit (Applied Biosystems, Cat. No. 4368814). The relative gene expression was measured according to 2^-ΔΔCT^ methods. The house keeping gene 36B4 was used for internal control. The Primer sequences were: 36B4, F: GGC GAC CTG GAA GTC CAA CT; R: CCA TCA GCA CCA CAG CCT TC. KLK2, F:AGT CAT GGA TGG GCA CAC TG, R:GCC AGA CCT GGC TAT TCT. KLK3, F:GAC CAA GTT CAT GCT GTG TGC, R:CCA CTC ACC TTT CCC CTC AAG. TMPRSS2, F:GTG AAA GCG GGT GTG AGG AG, R:CTG GTG ACC CTG AGT TC. FKBP5, F:CTC CCT AAA ATT CCC TCG AAT GC, R:CCC TCT CCT TTC CGT TTG GTT. AR, F:GAC ATG CGT TTG GAG ACT GC, R: TTT CTT CAG CTT CCG GGC TC, TEAD1,F: ATG GAA AGG ATG AGT GAC TCT GC, R: TCC CAC ATG GTG GAT AGA TAG C, TEAD4, F: GAA CGG GGA CCC TCC AAT G, R: GCG AGC ATA CTC TGT CTC AAC

### ChIP qPCR

ChIP (Chromatin Immuno-precipitation) assay was performed as previously study description (Reddy *et al*, 2009), In brief, cells were cross-linked by adding either formaldehyde to a final concentration of 1% for 10min or 2 mM DSG crosslinker (CovaChem, Cat. No.13301) at room temperature for 1 h followed by secondary fixation with 1% formaldehyde (Pierce, Cat. No. 28908) for 10 min and quenched by adding glycine. Subsequently, cells were washed with cold PBS and subject to cell lysis. The cell extracts were subject for sonication. After centrifuge, the cell extracts were incubated with prepared AR/TEAD antibody-Dynabeads for 1 hour room temperature and another 1 hour at 4℃, washed in wash buffer for 5 times and de-cross-linked ChIP in elution buffer at 65℃ overnight. The antibodies used in ChIP-qPCR were anti-AR (Sigma, Cat. No. A9853), anti-TEAD4 (Santa Cruz, Cat. No. sc-101184), and anti-PanTEAD (Cell signaling Technology, Cat:13295). Anti-rabbit IgG dynabeads (Invitrogen, Cat: 11204D) and anti-mouse IgG dynabeads (Invitrogen, Cat. No. 11031) were used to bind antibodies. The enriched DNA was extracted via DNA extraction kits (Qiagen, Cat. No. 28106) and subject to quantitative PCR analysis. The Primer sequences for ChIP-qPCR were: KLK2, F: CCA TCT TGC AAG GCT ATC TGC TG, R: TGT GTC TTC TGA GCA AAG GCA AT; KLK3, F: ATA CTG GGA CAA CTT GCA AAC CT, R: CAG GCT TGC TTA CTG TCC TAG ATA A; FKBP5, F: GGA GCC TCT TTC TCA GTT TTG, R: CAA TCG GAG TGT AAC CAC ATC; TMPRSS2, F: GTG GCC CAC TTC CTC, R: CAC ACA GCA AGG CAG AGG ACA; CTGF, F: TGT GCC AGC TTT TTC AGA CG, R:TGA GCT GAA TGG AGT CCT ACA CA.

### ChIP-seq experiment and data analysis

ChIP-Seq libraries were generated using KAPA HTP Library Preparation Kit Illumina® platforms (KAPA, KR0426) according to manufacturer’s protocol. Briefly, the ChIP DNA were ligated with multiplex adapters and amplified by PCR to generate libraries, after purification and size selection, the libraries were checked on the Agilent 2100 Bioanalyzer. After samples were quantified, normalized and pooled, final samples run on the Illumina HiSeq 2500 using the 75bp high output sequencing kit for single-end sequencing at UTSW next generation sequencing core. Reads were trimmed and aligned to the human genome (hg19) using ‘Bowtie2’, reads were sorted by ‘Samtools’ subsequently. Peaks were identified using MACS2 v2.1.1 with the p-value cutoff 1e-5. After peak calling, ENCODE human blacklisted regions were removed from the peak files subsequently. Peak overlapping analysis, motif enrichment analysis and peak annotation were performed using ‘HOMER’ v4.9. ChIP-seq signal tracks were generated by ‘DeepTools’ and normalized by RPKM. ChIP-seq data are deposited in the Gene Expression Omnibus (GEO) database (Assessing number: GSE208606). Signal tracks were visualized by Integrative Genomics Viewer (IGV) software. Heatmaps and signal plots for the ChIP-seq peak subsets were generated using ‘DeepTools’.

### RNA sequencing and data analysis

The global gene expression analysis (Vehicle vs XMU-MP-1 treated groups) was based on RNA sequencing platform from BGI (Beijing Genomic Institute). Cellular RNA was extracted using Qiagen RNA extraction kit (Qiagen; Cat: 74104) according to the manufacturer’s instructions. The cellular RNA was sent to BGI Genomics (https://www.bgi.com) for RNA sequencing. RNA was quality-accessed with an Agilent 2100 Bioanalyzer (Agilent RNA 6000 Nano Kit) with RNA integrity number above 9 for library construction. The total RNA was used for library construction according to the protocol of BGISEQ-500 platform. The libraries were sequenced using BGISEQ-500 platform. Then the FASTQ sequencing files were aligned to the hg19 human genome using STAR aligner with uniquely mapped reads kept for further analysis. The RNA sequence data are deposited in the Gene Expression Omnibus (GEO) database (Assessing number: GSE186177). Pathway analysis of differentially expressed genes (DEGs) (FPKM>5, P value < 0.05 and Fold change >1.5) was performed using Metascape (https://metascape.org) and heatmap was plotted by http://www.bioinformatics.com.cn, a free online platform for data analysis and visualization. For gene set enrichment analysis of RNA-seq data, gene sets of Hallmark Androgen Response and Cordenonsi YAP Conserved Signature were used and downloaded from Molecular Signatures Database v7.4, GSEA was implemented using the GSEA 4.1.0 software, with default parameters. Volcano plot was generated using ‘ggplot2’ package in R (threshold P<0.05 and fold change>1.5).

### Analysis of TCGA prostate cancer data sets

Expression analysis of *YAP, TAZ, AMOTL2*, *KLK3 and KLK2* in prostate cancer tissues and normal tissues were performed by http://ualcan.path.uab.edu/ online tool. For the correlation analysis between AR and YAP target genes, MORPHEUS (https://software.broadinstitute.org/morphe us/) was used for generation of gene correlation heatmap, spearman correlation analysis between genes were performed by cBioprotal (http://www.cbioportal.org/) used 489 samples of Prostate Adenocarcinoma (TCGA, PanCancer Atlas).

### Cell proliferation assay

LNCaP, C4-2, 22RV1 and R1-D567 cells were transfected with siYAP, siTEAD1/3/4, siLATS1/2 or scramble siRNAs in 24-well plates. 24 hours after transfection, the cells number was counted, and 10000 cells were seeded into 96-well plates. For drug treatment assays, 2000-10000 cells were treated with 1-5 μM XUM-MP-1 or 20 μM Enzalutamide in 96-well plates. The relative cell viability was measured at indicated time points. Cell numbers were determined using the WST-1 cell proliferation reagent (Sigma Aldrich, Cat. No. 5015944001).

### Xenograft tumor models

The procedures for all animal experiments were reviewed and approved by the IACUC of UT Southwestern Medical School. Six-week-old male NOD scid gamma (NSG) mice were used, and tumor cell were implanted subcutaneously into the dorsal flank of the mice(1 × 10^6^ cells suspended in 100ul PBS with 50% Matrigel). When tumor xenografts reached a mean volume of ∼100 mm3 (length × width^2^ /2), mice were randomly assigned to experimental treatment groups (6-8 mice/ group). XMU-MP-1 (dissolved in 5% dextrose in water) was administered intraperitoneally daily with 3-15mg/kg body weight for indicated time, and the control group were injected with the solvent. Dox in PBS was injected intraperitoneally with 20mg/kg body weight every day. Tumor size was measured using digital calipers. At the end of the studies mice were sacrificed, and liver, spleen and tumors were harvested and weighed.

## Statistics

Student’s t-test was used for comparisons. A P-value of < 0.05 is considered significant. Error bars on the graphs were presented as the s.d.

## ACKNOWLEDGEMENTS

We thank Drs. Ping Mu and Ganesh Raj for reagents. The work was supported by grants from NIH to J.J. (R35GM118063) and Y.Y (R01CA222571), and Welch Foundation (I-1603) to J.J, J.J. is a Eugene McDermott Endowed Scholar in Biomedical Science at UTSW.

## Author contributions

J.Z. and J.J. conceived the study. J.J, J.Z, S.Z., and X. L. designed the experiments. S.Z., X. L., Y.C., Y.L., and Y.Y. performed the investigation and validated the results. J.J. wrote the manuscript. J.J. and Y.Y. acquired the funding.

## Extended view figure legends

**Fig. EV1.**
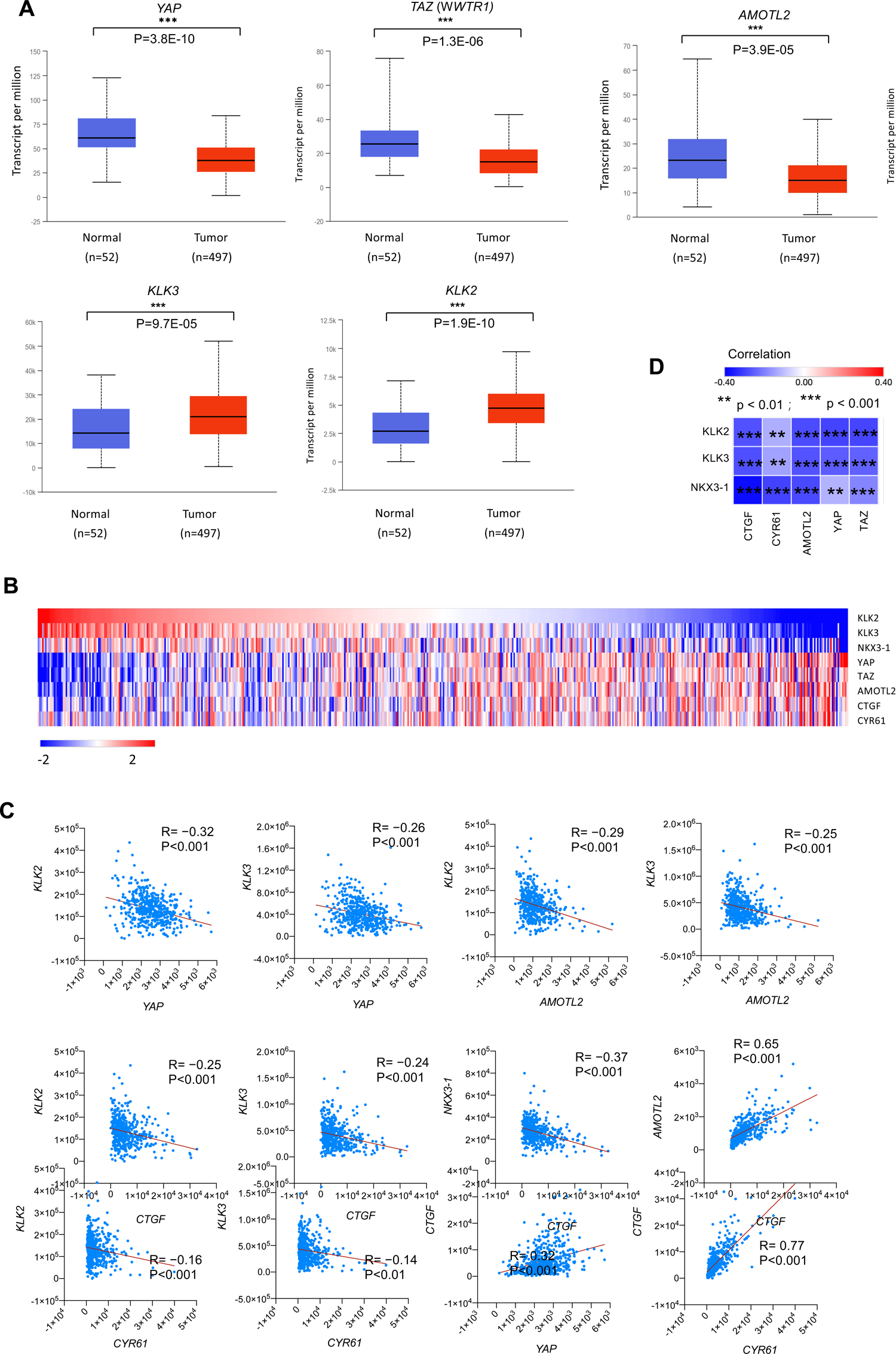
Reverse correlation between AR and YAP signaling activity in prostate cancer. (A) Analysis of TCGA database shows that mRNA levels for *YAP*, *TAZ* and *AMOTL2* are lower whereas those for *KLK2* and *KLK3* are higher in prostate cancer samples (n=497) compared with normal prostate tissues (n=52). (B) Heatmap for the expression of AR signaling and YAP signaling pathway target genes in TCGA prostate cancer data sets (in the diagram, red represents high gene expression levels, blue represents low gene expression levels). (C-D) The correlation between AR signaling and YAP signaling in TCGA prostate cancer data sets. In the heatmap diagram (D), the horizontal and vertical coordinates represent genes, and different colors represent correlation coefficients (red represents positive correlation, blue represents negative correlation), the darker the color represents the stronger the correlation.

**Fig. EV2.**
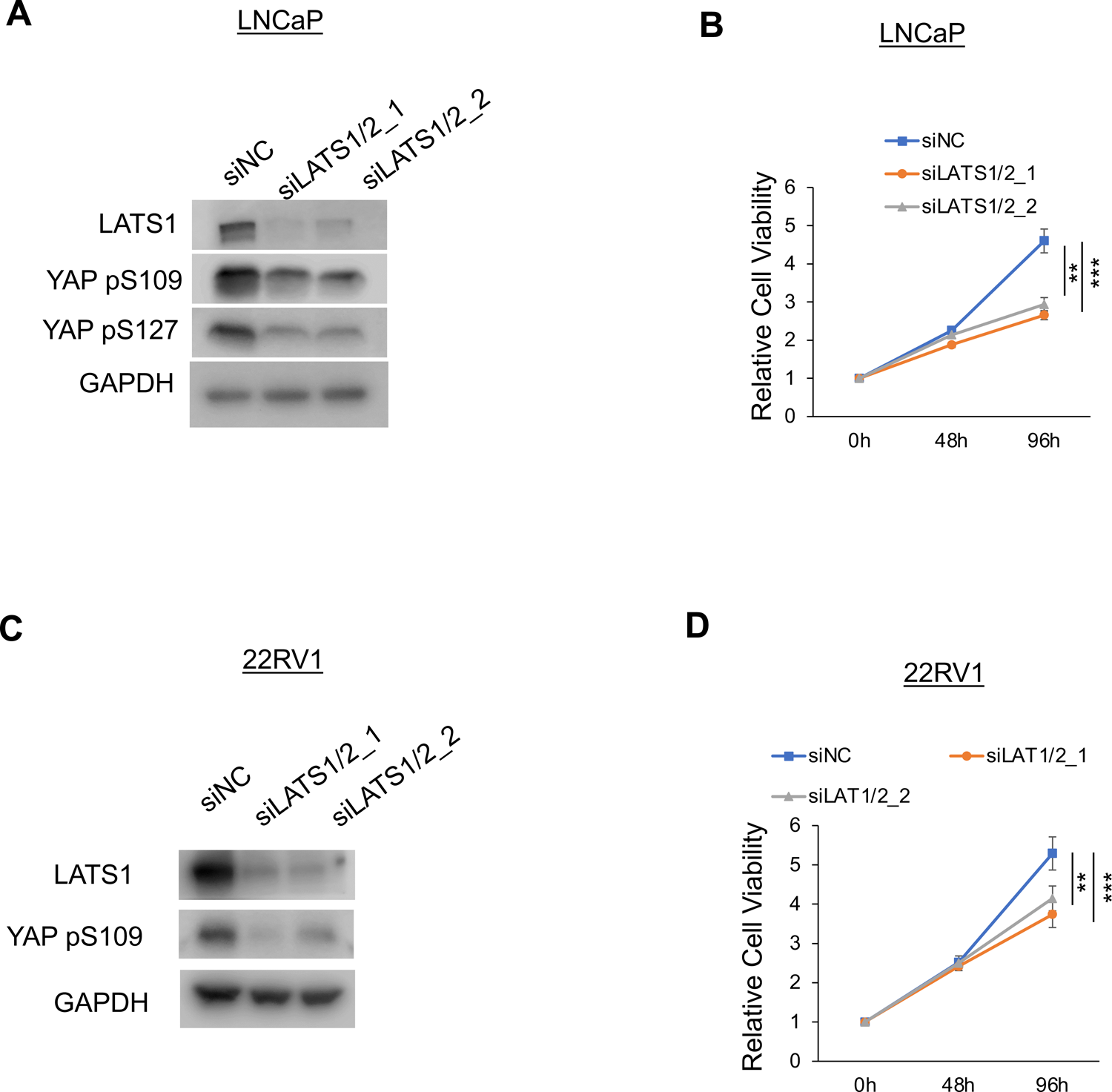
LATS1/2 knockdown inhibited LnCaP and 22RV1 cell growth *in vitro*. (A, C) Western blot analysis of LATS1 and YAP phosphorylation in LNCaP (A) or 22RV1 (C) cells treated with the control or LATS1/2 siRNA. (B, D) Relative growth of LNCaP (A) or 22RV1 (C) cells treated with the control or LATS1/2 siRNA. Results are representatives for two independent experiments. Data are ± SD. **P<0.01, ***P<0.001 (Student’s t-test).

**Fig. EV3.**
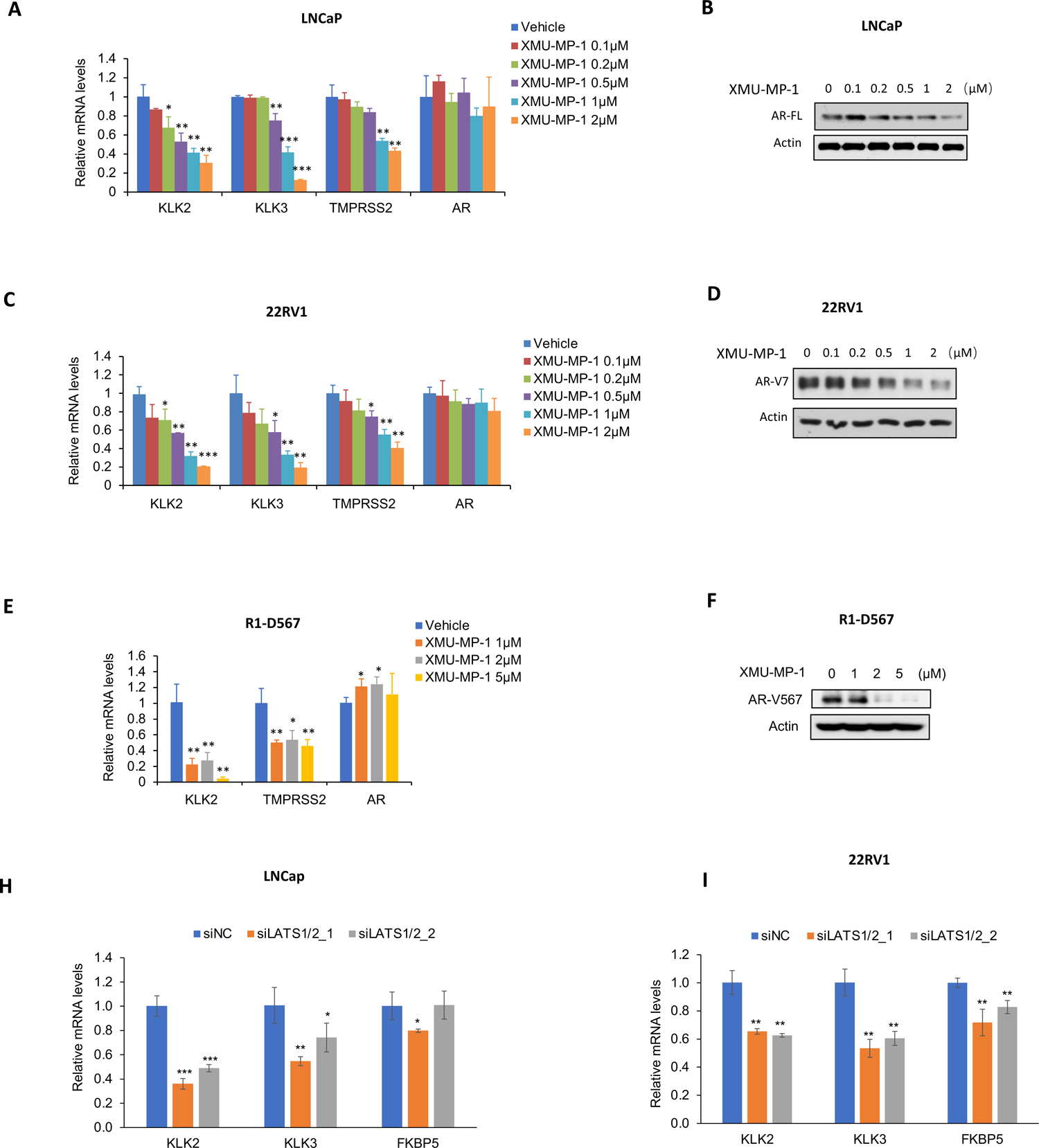
XMU-MP-1 inhibited AR target gene expression. (A-F) Relative mRNA levels of AR and the indicated AR target genes (A, C, E) and western blot analysis of AR and YAP in LNCaP (A-B), 22RV1 (C-D), or R1-D567 cells treated with vehicle or XMU-MP-1 at the indicated concentrations for 8 hours. XMU-MP-1 inhibited the expression of AR target genes in a dose dependent manner. (H-I) Relative mRNA levels of AR target genes in LNCap (H) or 22RV1 (I) cells expressing control or LATS1/2 siRNA. Results are representatives for two independent experiments. Data are ± SD. *P<0.05, **P<0.01, ***P<0.001 (Student’s t-test).

**Fig. EV4.**
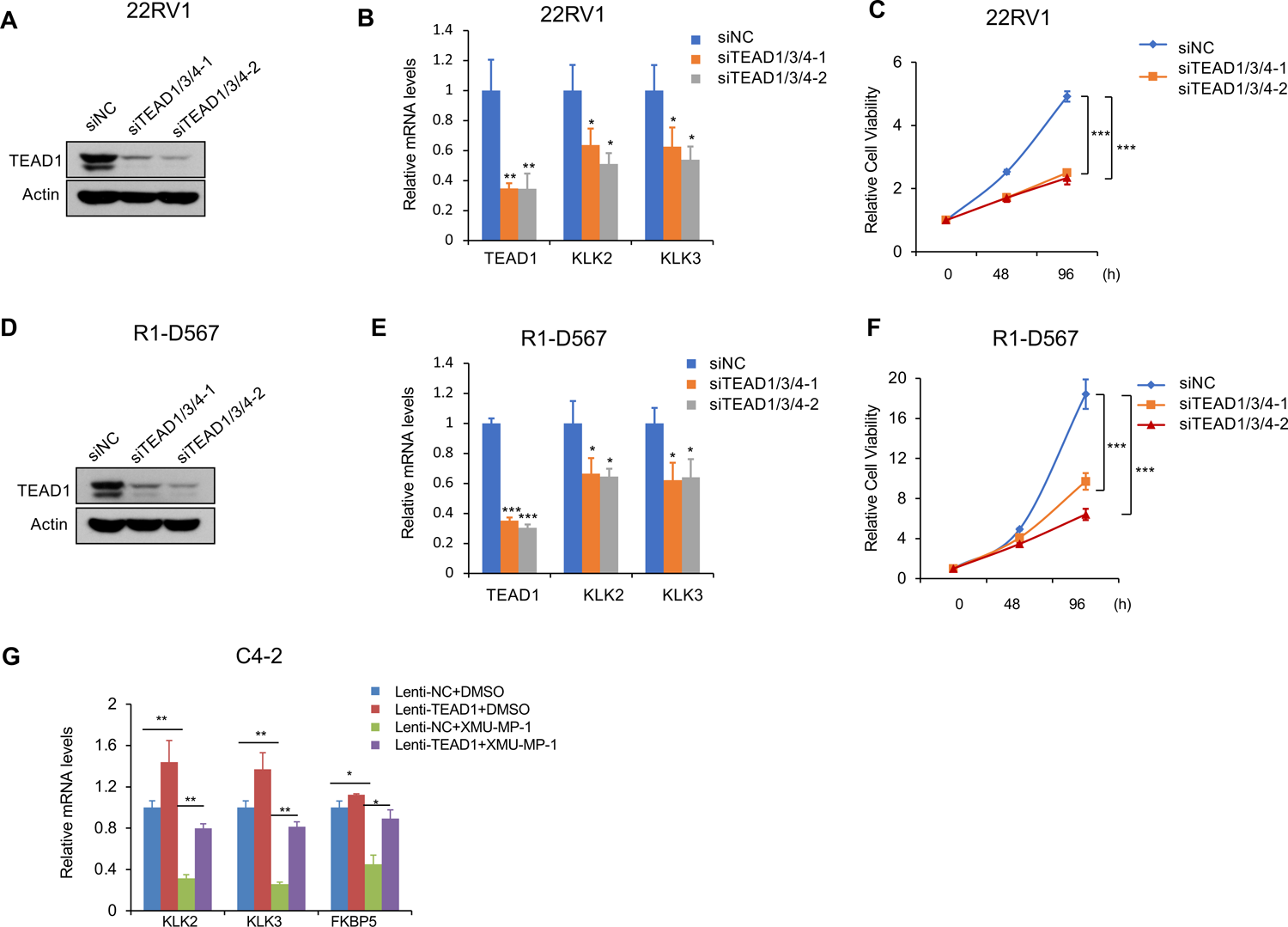
TEAD knockdown affected AR target gene expression and 22RV1 and R-D567 PCa growth. (A-F) Western blot analysis of TEAD1 (A, D), relative mRNA levels of *TEAD1*, *KLK2* and *KLK3*. (B, E), and relative growth (C, F) of 22RV1 (A-C) or R1-D567 (D-F) PCa cells treated control or two independent TEAD1/3/4 siRNA (siTEAD1/3/4_1 or siTEAD1/3/4_2). (G) Overexpression of TEAD1 partially rescued the expression of indicated AR target genes inhibited by XMU-MP-1. Results are representatives for three independent experiments. Data are ± SD. *P<0.05, **P<0.01, ***P<0.001 (Student’s t-test).

**Fig. EV5.**
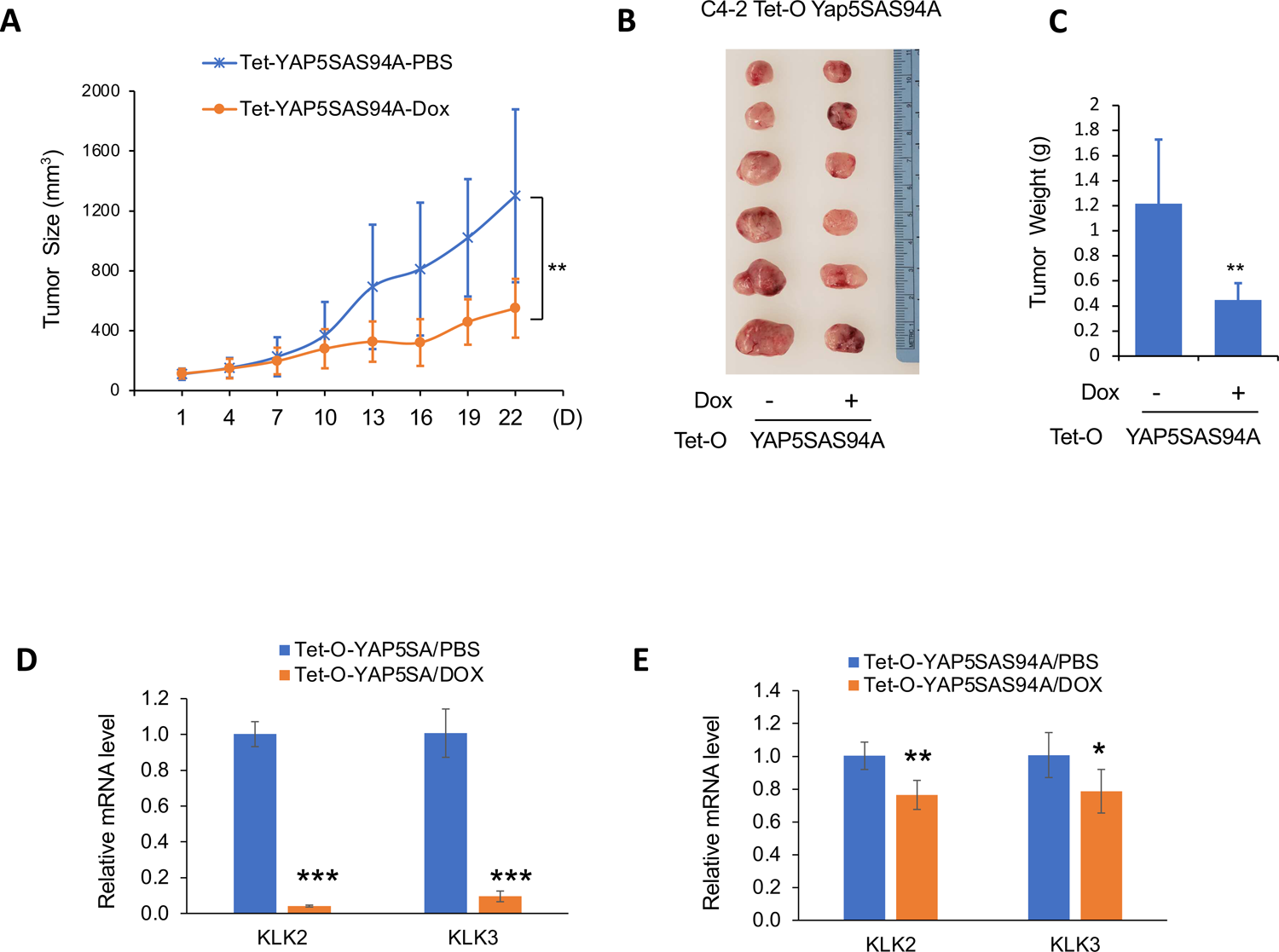
The effect of YAP-5SAS94A overexpression on C4-2 PCa growth *in vivo*. (A-C) Male NOD scid gamma (NSG) mice bearing C4-2 tumors that express Tet-O-YAP-5SAS94A were injected i.p. with PBS containing Dox (20mg/kg) or PBS daily for the indicated period of time. Tumor growth curve (A), photograph of tumor samples (B), and quantification of tumor weight (C) at the end of treatment were shown. (D-E) Relative mRNA levels of *KLK2* and *KLK3* in xenograft C4-2 tumors expressing express Tet-O-YAP-5SA (D) or Tet-O-YAP-5SAS94A (E) from mice treated with PBS or DOX for 30 days. Data are ± SD. *P<0.05, **P<0.01, ***P<0.001 (Student’s t-test).

**Fig. EV6.**
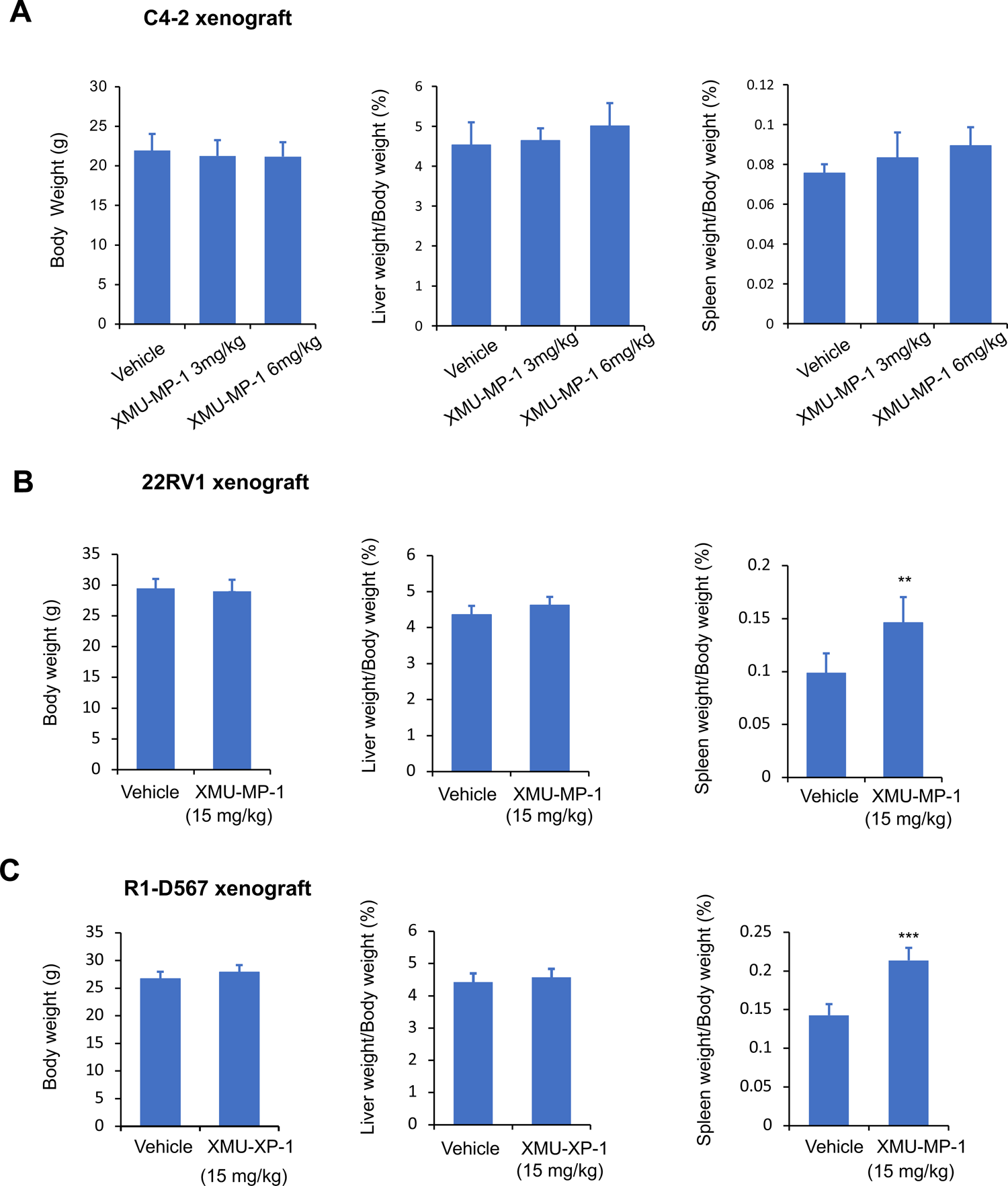
XUM-MP-1 was well-tolerated in mice. (A-C) Body weight (left), liver weight/body weight (middle) and spleen weight /body weight (right) of mice bearing C-42 (A), 22RV1 (B), or R1-D567 tumors and treated with daily XMU-MP-1 at the indicated concentrations for 25 days (A), 14 days (B), or 15 days (C). Data are ± SD. **P<0.01 (Student’s t-test).

**Fig. EV7.**
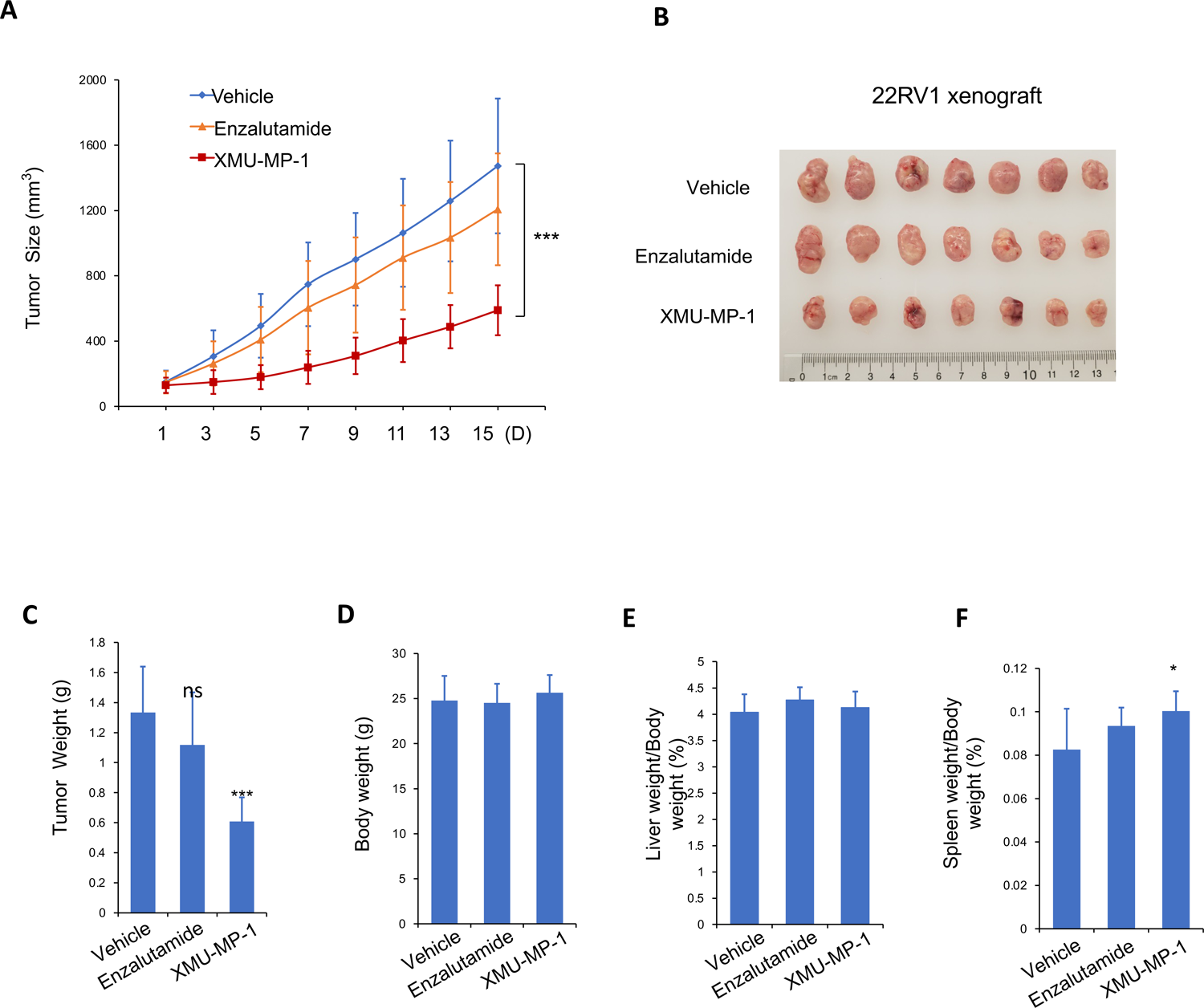
XMU-MP-1 but not Enzalutamide inhibited 22RV1 tumor growth in xenografts. (A-F) Male NOD scid gamma (NSG) mice bearing 22RV1 tumors were treated daily with XMU-MP-1 (15mg/kg), enzalutamide (10mg/kg) or vehicle for the indicated time. Tumor growth curve (A), photograph of tumor samples (B), tumor weight (C), body weight (D), liver weight/body weight (E) and spleen weight /body weight (F) at the end of treatment were shown. Data are ± SD. ns: not significant, *P<0.05, ***P<0.001 (Student’s t-test). n=7 mice for each group.

**Table EV1**

AR target genes downregulated by XMU-MP-1

**Table EV2**

AR/TEAD co-binding peaks in XMU-MP-1 downregulated AR target genes

